# Reconstructing True 3D Spatial Omics at Single-Cell Resolution

**DOI:** 10.64898/2026.04.28.721395

**Authors:** Yuhang Yang, Yiming Luo, Kai Zhang, Yonggan Bu, Zheng Xia, Haoxin Peng, Rui Yan, Qi Liu, Defu Lian, Yang Chen, Lin Shen, Enhong Chen

**Author notes:** These authors contributed equally to this work (Yuhang Yang, Yiming Luo).

## Abstract

Capturing the three-dimensional (3D) organization of cells is essential for deciphering complex biological processes, yet comprehensive 3D spatial omics is severely hindered by the destructive nature of physical sectioning and the depth limitations of intact tissue imaging. Current computational methods rely on 2.5D stacking of discrete slices, which inherently disrupts tissue topology and fails to resolve continuous depth-dependent molecular gradients. To bridge this gap, we introduce DeepSpatial, an Optimal Transport flow matching framework that models tissue evolution as a continuous dynamic vector field. By solving the underlying probability flow ODEs, DeepSpatial enables the direct extraction of uninterrupted, infinitely resolvable tissue states at arbitrary spatial depths. Using Deep STAR/RIBOmap 3D technologies, we demonstrate that DeepSpatial achieves improved 3D reconstruction fidelity relative to 2.5D approaches, yielding structures that more closely recapitulate native tissue microenvironments in real-world datasets. Across diverse spatial omics modalities, including spatial proteomics using imaging mass cytometry in human breast cancer and spatial transcriptomics using openST in head and neck squamous cell carcinoma metastatic lymph nodes, DeepSpatial produces biologically interpretable and high-fidelity reconstructions across datasets. We evaluated the scalability and robustness of DeepSpatial on a large-scale mouse brain dataset, reconstructing a continuous 3D cellular atlas comprising 39 million cells within 41.6 hours. Systematic downstream characterization validated its ability to recapitulate consistent spatial architectures, cell-type distributions, transcriptomic patterns, and microenvironmental structures across brain regions. Collectively, these results demonstrate DeepSpatial as a generalizable and efficient solution for true 3D spatial reconstruction across scales and modalities.

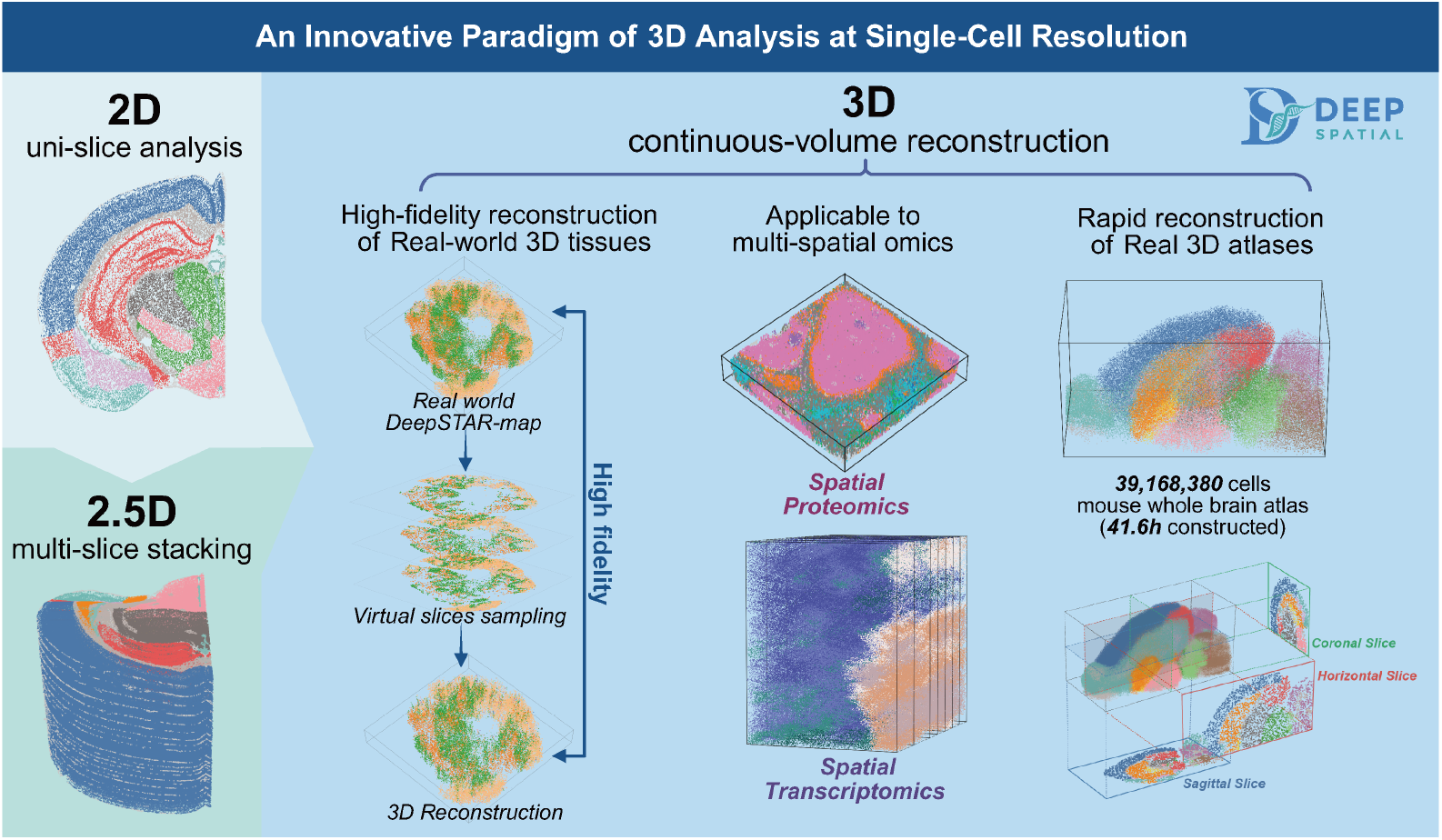

## 1 Introduction

Spatial omics has profoundly advanced our understanding of tissue architecture by enabling the mapping of gene expression patterns within their native spatial context^1–4^. Because the three-dimensional (3D) organization of cells is tightly coupled to their biological function, comprehensive 3D spatial profiling is essential for resolving complex biological systems^5,6^. Such approaches enable the reconstruction of continuous cellular and structural transitions, the delineation of spatial heterogeneity and molecular gradients, and the preservation of intact tissue architecture necessary to uncover localized transcriptional dysregulation^7–9^.

Current experimental approaches to capture 3D spatial omics generally fall into two categories: invasive serial sectioning and non-invasive intact thick-tissue profiling. Invasive techniques involve physically sectioning the tissue into sequential 2D slices, a process that inherently destroys the continuous 3D structure and introduces physical artifacts such as tears, folds, and geometric deformations^8,10,11^. Conversely, intact 3D imaging technologies attempt to profile thick tissue blocks without sectioning. However, these methods face severe physical and practical limitations: they are typically restricted to relatively thin tissue volumes (up to 200 µm), rendering organ-level 3D mapping impractical^12,13^. Additionally, these approaches are heavily constrained by resource-intensive, highly complex experimental protocols and prohibitive costs.

To overcome the experimental bottlenecks associated with physical sectioning and imaging depth, researchers have developed various computational frameworks to achieve 3D spatial analysis using existing 2D multi-slice datasets. Early computational approaches^14–17^primarily relied on spatial alignment and registration techniques. By applying rigid or non-rigid transformations to align discrete, consecutive 2D slices into a common 3D coordinate system, these methods constructed an initial spatial stack of the tissue. Building upon this foundation, recently developed generative methods^18–20^ have further advanced the field by moving beyond simple alignment to data imputation. By inferring and interpolating virtual slices between adjacent experimentally measured sections, these generative approaches effectively fill the inherent physical gaps caused by sparse sampling. Consequently, this virtual interpolation transforms the originally fragmented sequence into a denser stack, providing a more refined view of the tissue volume.

Despite these computational advancements, current frameworks fundamentally construct 2.5D models that rely on the spatial stacking of discrete 2D planes^18^. Even with the aid of dense virtual interpolation, the underlying representation remains inherently discontinuous and resolution-limited^21,22^. Excessive interpolation also introduces spurious spatial artifacts and redundant computations. This dimensional fragmentation between adjacent layers forcibly disrupts the continuous spatial connectivity of the tissue microenvironment, often obscuring subtle depth-dependent molecular gradients and blurring fine anatomical boundaries^23^. Such discrete frameworks struggle to accurately reconstruct the complex 3D cellular neighborhoods and topological structures that span multiple layers. To truly capture the uninterrupted topology and fluid transitions within biological microenvironments, transitioning from discrete 2.5D stacking to continuous, real 3D modeling is strictly necessary.

To address this fundamental limitation, we propose DeepSpatial, a generative framework based on flow matching^24,25^that seamlessly reconstructs the continuous 3D tissue microenvironment from discrete measured slices. DeepSpatial reframes the multi-modal transitions between physical slices as a mixed continuous-discrete probability density evolution. Methodologically, we first establish biologically rigorous cell-to-cell correspondences via Unbalanced Optimal Transport (UOT)^26^, explicitly constrained by spatial, transcriptomic, and categorical cell-type affinities. To model the underlying continuous dynamics, we introduce a scalable Transformer^27,28^architecture that employs a unified tokenization strategy to process heterogeneous cell states within a shared high-dimensional vector field. By solving the learned Probability Flow Ordinary Differential Equation (ODE)^29,30^with a density-preserving bidirectional sampling strategy, DeepSpatial enables the direct and exact recovery of uninterrupted tissue topology at any arbitrary physical depth. Consequently, DeepSpatial completely overcomes the structural fragmentation of physical sectioning, achieving a paradigm shift in spatial omics from slice-level 2.5D stacking to biologically faithful, volume-level 3D continuous modeling.

Comprehensive experimental validation across diverse biological systems, including tumor microenvironments, metastatic lymph nodes, and a 39-million-cell mouse whole-brain atlas, demonstrates that DeepSpatial achieves a transformative shift from discrete 2.5D stacking to biologically faithful 3D continuous modeling. To facilitate broad adoption and further development within the community, the DeepSpatial framework, along with its interactive documentation, full source code, and reproducibility tutorials, is publicly accessible at our Project Homepage and GitHub Repository.

## 2 Results

### 2.1 Overview of DeepSpatial

DeepSpatial is a generative deep-learning framework designed to construct a continuous, true 3D tissue atlas from sparsely sampled 2D spatial omics sections (Fig. 1). By reframing 3D reconstruction as a continuous probability density evolution, the model seamlessly bridges physical z-axis gaps. As illustrated in Fig. 1a, the framework processes multi-layer 2D spatial data by integrating spatial coordinates, high-dimensional omics features (genes and proteins), and physical slice metadata (thickness and interval). Through this continuous generative process, DeepSpatial systematically transforms these discrete inputs into a spatially continuous 3D virtual tissue microenvironment (Fig. 1b), wherein every synthesized cell is assigned exact 3D coordinates, omics expressions and cell-type identities.

**Fig. 1.**
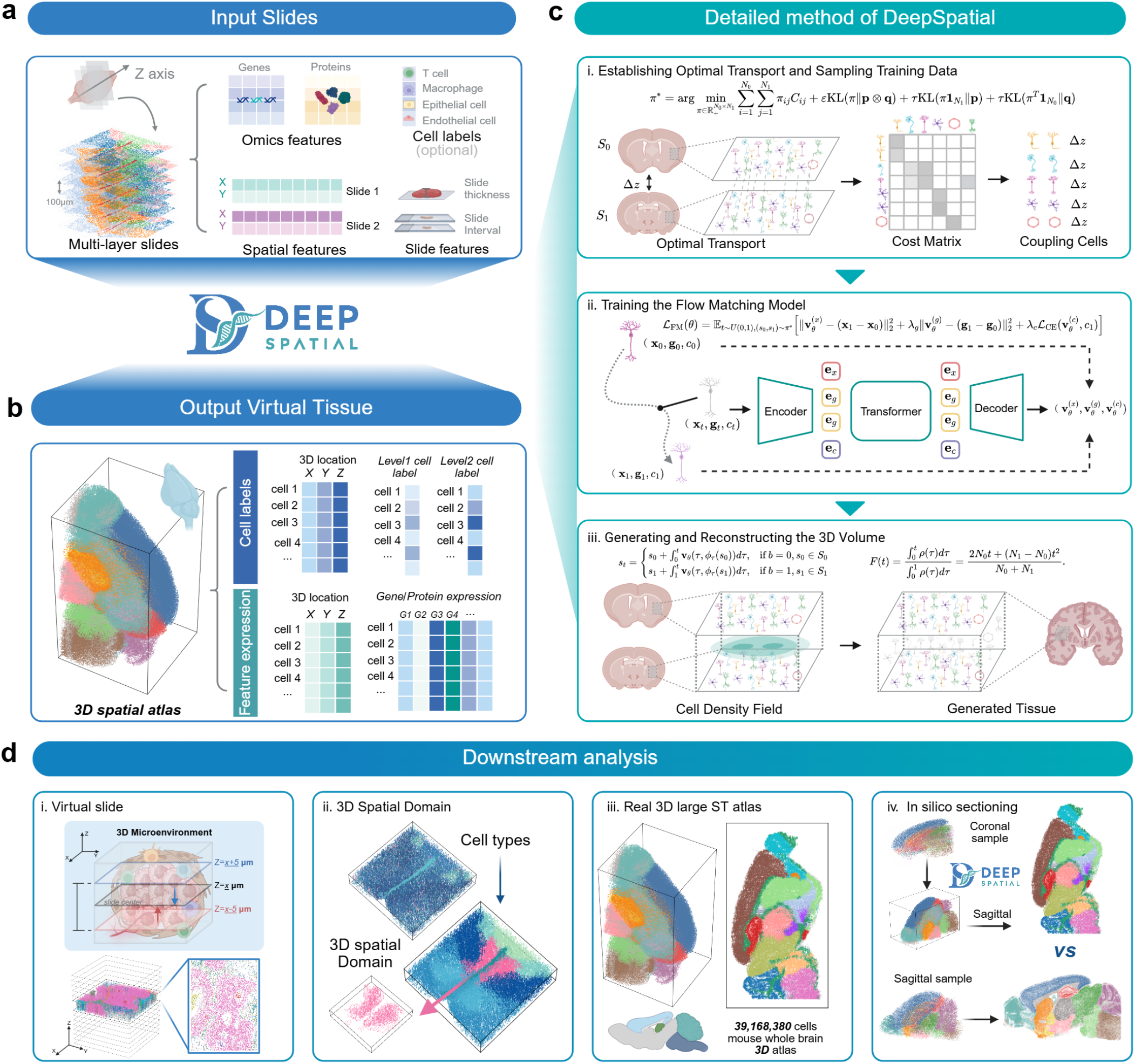
Overview of the DeepSpatial framework. **a**, The framework accepts discrete, multi-layer 2D spatial omics slices, incorporating spatial coordinates, omics expressions (genes and proteins), optional cell labels, and physical slice metadata (thickness and interval distances). **b**, DeepSpatial reconstructs a fully continuous 3D spatial atlas, assigning precise 3D spatial coordinates, cell-type identities, and high-dimensional omics expressions to each generated virtual cell. **c**, The generative process consists of three key stages: (i) aligning cells across adjacent slices via Unbalanced Optimal Transport (UOT) based on spatial and transcriptomic affinities; (ii) learning a continuous cell-state transition vector field with a multi-modal Transformer via flow matching; and (iii) generating the 3D volume by solving Probability Flow ODEs guided by interpolated cell density. **d**, The 3D microenvironment enables advanced spatial analyses, including the extraction of precise virtual slices at any arbitrary depth (i), the identification of topologically complex 3D spatial domains (ii), the reconstruction of large organ-scale atlases (iii), and flexible multi-planar *in silico* sectioning across sagittal or coronal axes (iv).

The computational core of DeepSpatial models the multi-modal transitions between physical slices as a mixed continuous-discrete probability density evolution (Fig. 1c). First, to establish biologically rigorous alignments across physical gaps, the model computes cell-to-cell correspondences via UOT. This is governed by a cost matrix that balances spatial proximity and transcriptomic similarity while accounting for cellular proliferation or apoptosis. Second, we employ a scalable multi-modal Transformer trained via flow matching to learn a shared, continuous vector field representing the dynamic evolution of heterogeneous cell states. Finally, by solving the underlying Probability Flow ODE guided by an interpolated macroscopic cell density field, DeepSpatial enables the direct and exact synthesis of tissue states at any arbitrary spatial depth.

This paradigm shift from discrete 2.5D stacking to continuous 3D modeling unlocks comprehensive downstream analytical capabilities (Fig. 1d). The uninterrupted 3D microenvironment allows for the extraction of highly resolved virtual slices at arbitrary depths and the precise delineation of topologically complex 3D spatial domains. Furthermore, the framework’s scalability supports the reconstruction of large, organ-level spatial architectures, such as a 39-million-cell mouse whole brain atlas, and enables flexible multi-planar in silico sectioning across sagittal or coronal axes.

### 2.2 Benchmarking DeepSpatial against real 3D spatial data

To further evaluate the spatial reconstruction fidelity of DeepSpatial, we tested it on three real 3D datasets: Deep-STARmap (cSCC and mouse brain) and Deep-RIBOmap (mouse brain), each derived from 150 *µ*m continuous tissue volumes^13^. We first downsampled the ground-truth 3D data into sparse pseudo-2D sections (Fig. 2b), then reconstructed full volumes using DeepSpatial (true 3D) and SpatialZ (2.5D stacking), respectively. Finally, we quantitatively compared the two reconstructions against the ground truth at both spatial and transcriptomic levels (Fig. 2a).

**Fig. 2.**
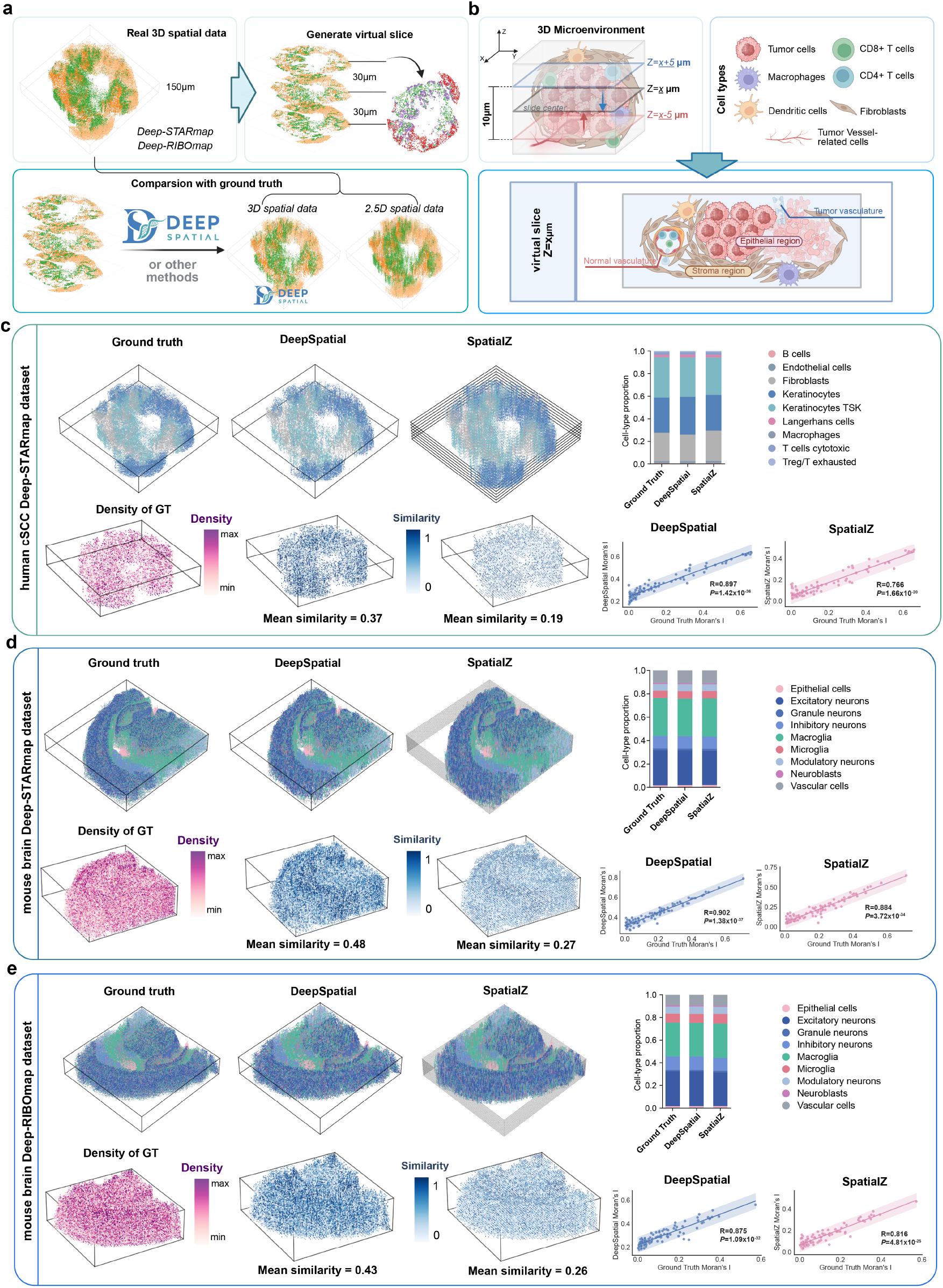
Performance evaluation of DeepSpatial for 3D spatial tissue reconstruction. **a**, Schematic benchmarking workflow using real 3D spatial transcriptomic data. **b**, Simulated 3D tissue model for validation. **c–e**, Quantitative evaluation on human cSCC Deep-STARmap (c), mouse brain Deep-STARmap (d), and mouse brain Deep-RIBOmap (e). 3D renderings of ground truth, DeepSpatial 3D reconstruction, and SpatialZ 2.5D stacking are shown alongside cell density, 3D cosine similarity, cell-type distribution, and Pearson correlation of gene-wise Moran’s I values.

We first confirmed that cell-type proportions were highly consistent among ground-truth, 3D-reconstructed, and 2.5D-imputed data. We then divided each 3D volume into local patches (50 ×50 × 30 for cSCC; 50 × 50 × 50 for mouse brain) and computed the cosine similarity of cell-type distributions within each patch. DeepSpatial-reconstructed tissues showed significantly higher similarity to ground truth: 0.37 (cSCC), 0.48 (STARmap), 0.43 (RIBOmap), whereas 2.5D imputation only achieved 0.19, 0.27, and 0.26 (Figs. 2c–e), which is attributed to severe discontinuity along the z-axis. We further validated that virtual cross-sections from our 3D reconstruction closely matched the real data (Figs. S1b, S2b, S3b).

We next evaluated transcriptomic spatial structure by computing Moran’s I for the top 100 highly variable genes. Both reconstructions correlated well with ground truth, but DeepSpatial 3D consistently achieved higher accuracy: cSCC (*R* = 0.897 vs 0.766), mouse brain STARmap (*R* = 0.902 vs 0.884), RIBOmap (*R* = 0.875 vs 0.816) (Figs. 2c–e). Cell-type-specific marker genes were faithfully preserved in 3D reconstructions, including *CD3E* (T cells), *CD79A* (B cells), *Gad1*/*Gad2* (inhibitory neurons), and *Slc17a7* (excitatory neurons), confirming high transcriptomic fidelity.

### 2.3 DeepSpatial reveals immune–vascular organization in the breast cancer microenvironment

In real-world applications, spatial datasets typically consist of multiple physical sections acquired at fixed intervals. To validate DeepSpatial under such conditions, we applied it to a 15-slice imaging mass cytometry (IMC) dataset of breast cancer (BRCA), with 2 *µ*m thickness and 20 *µ*m inter-slice gaps (Fig. 3a)^11^. Independent clustering and annotation across slices confirmed consistent cell-type identities and marker expression prior to reconstruction (Figs. S4–S6). Using DeepSpatial, we reconstructed a continuous 3D tumor microenvironment spanning 280 *µ*m along the z-axis (Fig. 3a).

**Fig. 3.**
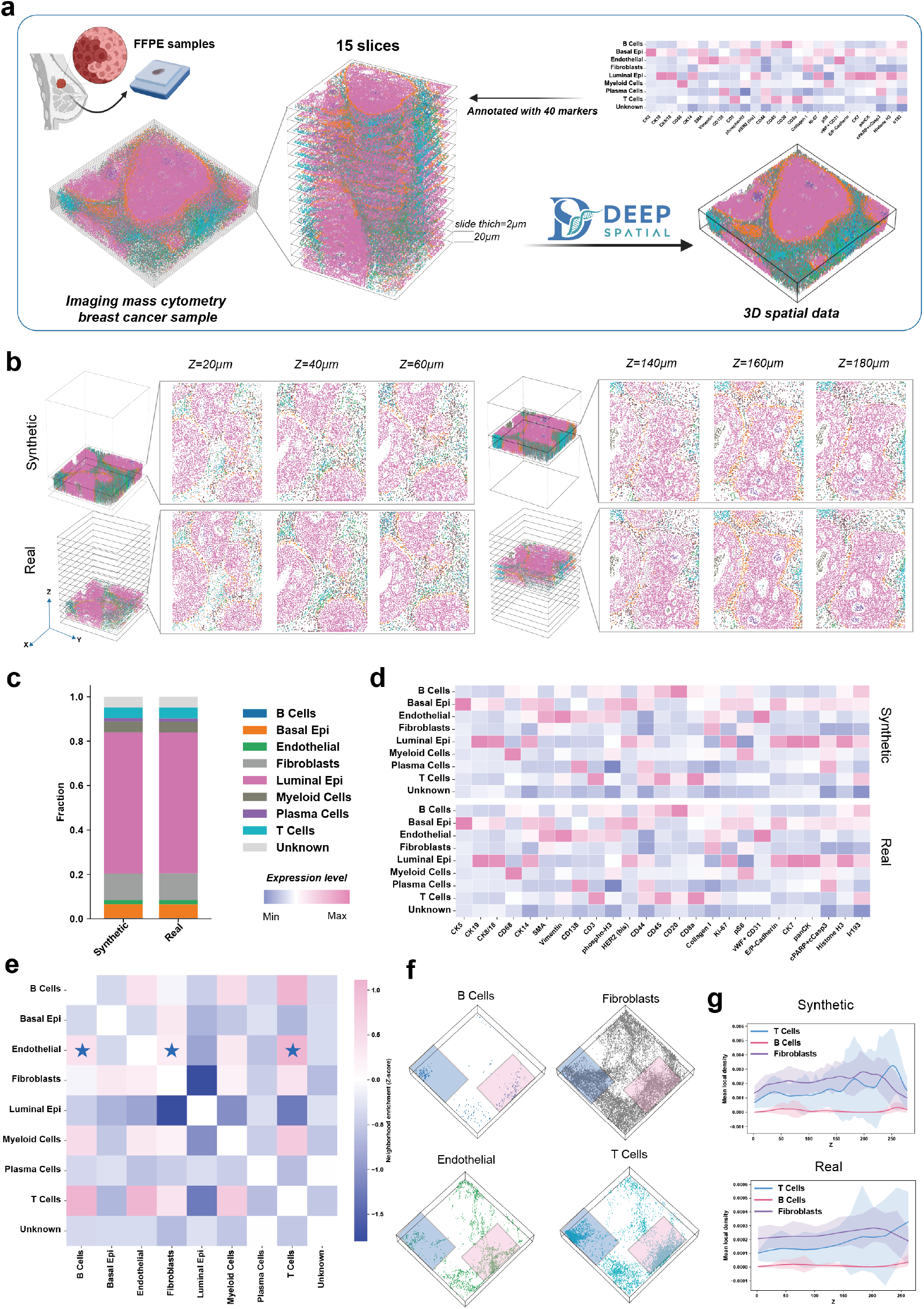
Application of DeepSpatial to BRCA IMC dataset. **a**, Workflow for 3D reconstruction of 15-slice IMC data. **b**, Comparison between real IMC slices and DeepSpatial-generated virtual slices. **c**, Cell-type proportion consistency between original and reconstructed tissue. **d**, Protein marker expression in real vs synthetic slices. **e**, Cellular neighborhood network highlighting endothelial-immune interactions. **f**, The 3D distribution of B Cells, fibroblasts, Endothelial Cells, and T Cells, and the blue, red areas show the co-location of different cell types. **g**, The Z-axis distribution of T Cells, B Cells, and Fibroblasts surrounding Endothelial cells.

Virtual slicing analysis showed that synthetic slices from the 3D volume closely matched the original IMC data, with fine-scale tissue architecture faithfully preserved (Fig. 3b). Sagittal and coronal virtual cross-sections further revealed that DeepSpatial effectively eliminated gaps from physical sectioning, yielding continuous tissue structures with clear epithelial, basal-like epithelial, and stromal compartments (Figs. S8a–d). Cell-type proportions were highly consistent between original and reconstructed data (Fig. 3c). At the protein level, cell-type-specific markers, including CD20 (B cells), CD3 (T cells), and CK5 (basal epithelial cells), exhibited nearly identical spatial patterns (Fig. 3d).

Vascular architecture is a key feature of the BRCA tumor microenvironment^31,32^. While original slices only show fragmented vessels, the full 3D reconstruction revealed a continuous, hierarchical vascular network with fine distal branches (Figs. S7c–e). Cell neighborhood analysis demonstrated that endothelial cells were significantly co-localized with T cells, B cells, and fibroblasts, indicating tight immune–vascular coupling (Fig. 3e). Spatial distribution along the z-axis further revealed that immune cells accumulated at 150–250 *µ*m, while endothelial cells were enriched near 50 *µ*m (Figs. S7f,g). Notably, immune clusters preferentially localized to distal vascular ends, revealing a spatially organized immune–vascular niche in BRCA (Fig. S7c).

### 2.4 3D Spatial Reconstruction of openST spatial transcriptomics dataset Using DeepSpatial

Spatial transcriptomics has recently emerged as a powerful tool for studying disease and tumor microenvironments. The openST technology enables whole-transcriptome spatial profiling and supports 3D spatial characterization through multiple tissue sections^33^. Here, we applied the DeepSpatial framework to reconstruct and complete 3D spatial structures from multi-layer openST data of metastatic lymph nodes in head and neck squamous cell carcinoma (HNSCC) (Fig. 4a).

**Fig. 4.**
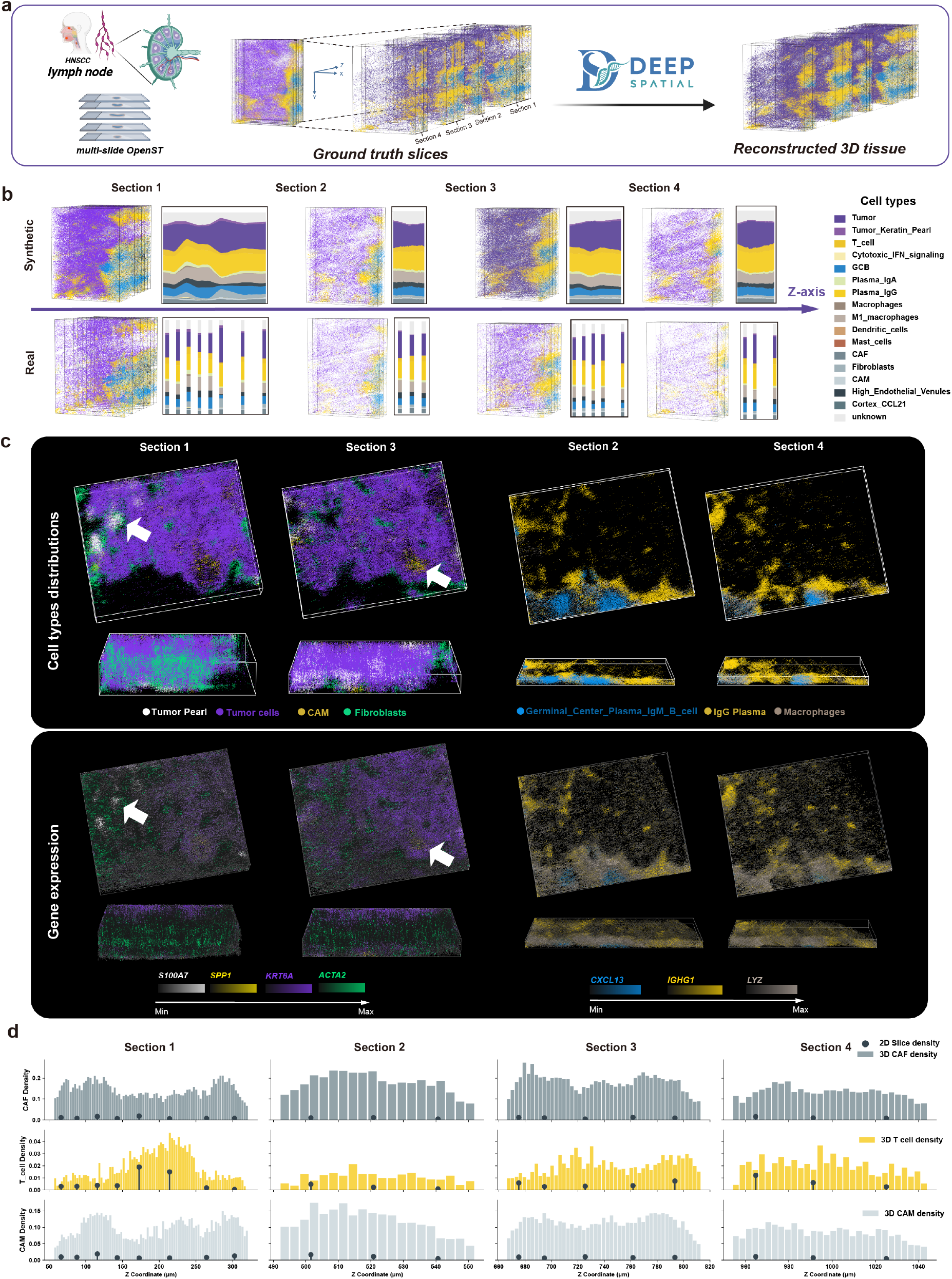
3D reconstruction of HNSCC metastatic lymph node openST data using DeepSpatial. **a**, Schematic reconstruction workflow. **b**, Cell-type distributions along z-axis: original (bottom) vs reconstructed (top). **c**, Cell-type distribution (top) and canonical marker expression (bottom), including *S100A7, SPP1, KRT6A, ACTA2, CXCL13, IGHG1, LYZ*. **d**, The 3D density distribution and 2D density distribution of CAF, T cells, and CAM surrounding Tumor cells.

Due to large and irregular inter-slice spacing that cannot support continuous whole-volume reconstruction, we split the dataset into four subgroups for independent 3D reconstruction (Figs. 4a–c, S5a,b). To avoid edge artifacts, all analyses were restricted to the maximal common region shared across all sections (Figs. 4b,c). DeepSpatial generated smooth cellular transitions along the z-axis, with cell-type proportions varying continuously rather than discretely (Fig. 4b). At the transcriptomic level, despite the high dimensionality and inherent noise characteristic of such data, gene expression patterns were recovered with high fidelity. Canonical, biologically relevant marker genes were consistently preserved across cell types, including *S100A7* in keratinized tumor cells, and *SPP1* in CAM populations (Figs. 4c, S10).

Cellular neighborhood analysis is a central component of spatial profiling, particularly in the metastatic tumor microenvironment (TME), where the spatial relationships between tumor, stromal, and immune cells are of primary interest. We characterized stromal and immune infiltration surrounding tumor cells in metastatic lymph nodes by quantifying local cellular densities within a 20 *µ*m radius of tumor cells (see Methods). Under comparable global cellular densities, volumetric density estimates in 3D tissues exceeded those derived from 2D sections and revealed more refined clustering patterns (Fig. 4e). In certain regions, 2D neighborhood analyses from individual sections were consistent with 3D observations. For example, in Section 1, T cells exhibited enrichment at depths of 200–230 *µ*m, whereas CAFs and CAMs were enriched in adjacent regions (Fig. 4e). However, in other regions, such as 700–740 *µ*m in Section 3, T cell clustering was not directly detectable in individual 2D sections (Fig. 4e). This discrepancy arises because cellular distributions across adjacent sections collectively contribute to enrichment patterns in 3D space, whereas 2D analyses neglect inter-slice information and thus underestimate the underlying spatial organization.

To more precisely characterize the tumor microenvironment, we further applied CellCharter^34^, a spatial domain identification method at single-cell resolution, to delineate diverse spatial architectures within metastatic lymph nodes. Tumor regions were spatially segregated from T cell–enriched areas by medullary cords and fibrotic capsules, with CAFs forming a barrier between stromal and tumor compartments (Figs. S12a–b). Notably, DeepSpatial accurately recapitulated both cellular composition and spatial architecture of each niche compared to original sections (Figs. S11a–c), demonstrating biologically interpretable high-fidelity reconstruction.

### 2.5 Scalable 3D reconstruction of a 39-million-cell mouse whole-brain spatial atlas

A major challenge in large-scale 3D tissue reconstruction is the high computational cost from high-dimensional omics features and millions of cells. We applied DeepSpatial to a CCF-aligned mouse whole-brain dataset with 129 sections (4,167,889 cells)^10^ (Fig. 5a), which expanded to 39,168,380 cells after 3D reconstruction. Notably, full reconstruction was completed within 41.6 hours on a mid-range server, demonstrating excellent efficiency and scalability (Table S3).

**Fig. 5.**
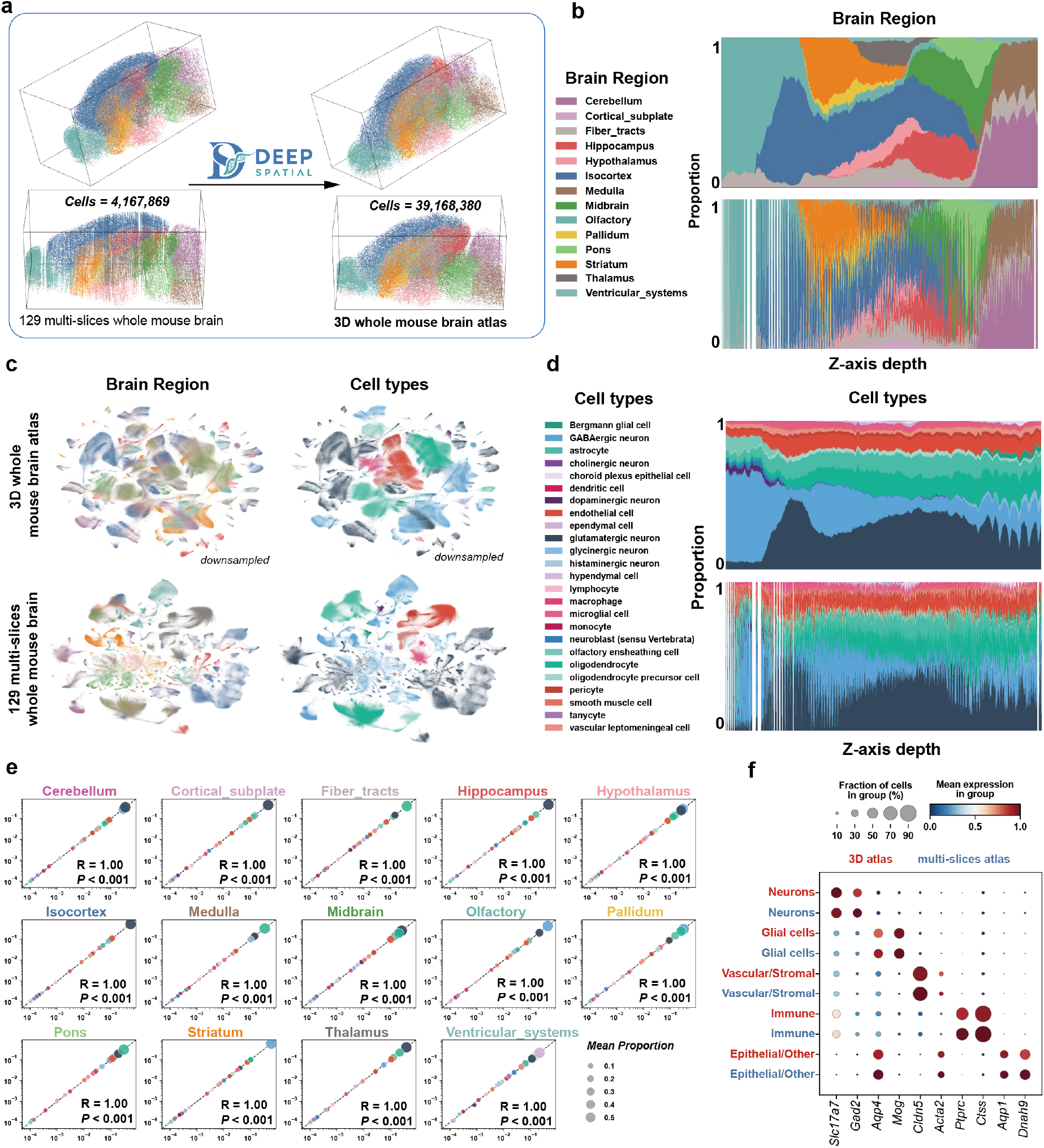
Scalable 3D reconstruction of a 39-million-cell mouse whole-brain atlas. **a**, Overview of 129-slice whole-brain reconstruction. **b**, Brain-region distribution along z-axis. **c**, UMAP comparison of brain regions and cell types. **d**, Cell-type distribution along z-axis. **e**, Spearman correlation of cell-type proportions per brain region. **f**, Gene expression consistency across major cell types.

We first examined cellular clustering in the 3D brain. UMAP embeddings showed similar cell-type and brain-region clustering between original and reconstructed data (Fig. 5c). A key advantage of true 3D reconstruction is z-axis continuity: relative to discontinuous raw sections, DeepSpatial produced smooth spatial distributions across x/y/z axes, especially along the discrete-sampled z-axis (Figs. 5b,d, S14a,b). Cell-type proportions within each brain region were strongly correlated between original and reconstructed tissues, confirming high-fidelity preservation of local structure (Fig. 5e).

Transcriptomic fidelity was maintained at the whole-brain scale. Major cell-type markers showed nearly identical expression levels and fractions: *Slc17a7* (excitatory neurons), *Cldn5* (vascular endothelial cells), *Ptprc* (immune cells) (Fig. 5f). Fine subpopulations were also preserved: *Gad2* (GABAergic neurons), *Hdc*/*Slc18a2* (histaminergic neurons), *Gja1*/*Gpr37l1* (astrocytes), *Sall1*/*Ctss*/*Mafb* (microglia), *Rgs5* (pericytes) (Fig. S13). These results demonstrate that DeepSpatial preserves cellular identity and transcriptome profile across millions of cells.

To further validate the biological authenticity of our 3D reconstruction, we performed multi-planar virtual slicing, a critical yet challenging task for 3D tissue modeling. We integrated whole-brain datasets from two mice for comprehensive comparison: Animal 1 (129 coronal slices) served as the ground-truth for interpolation, while Animal 2 (23 horizontal slices) provided independent validation for orthogonal slicing (Fig. 6a). In the coronal plane, virtual slices from the 3D-reconstructed tissue exhibited near-perfect concordance with the GT coronal sections, accurately recapitulating both brain region anatomy and spatial patterns of canonical markers including *Slc17a7* and *Gfap* (Fig. 6b).

**Fig. 6.**
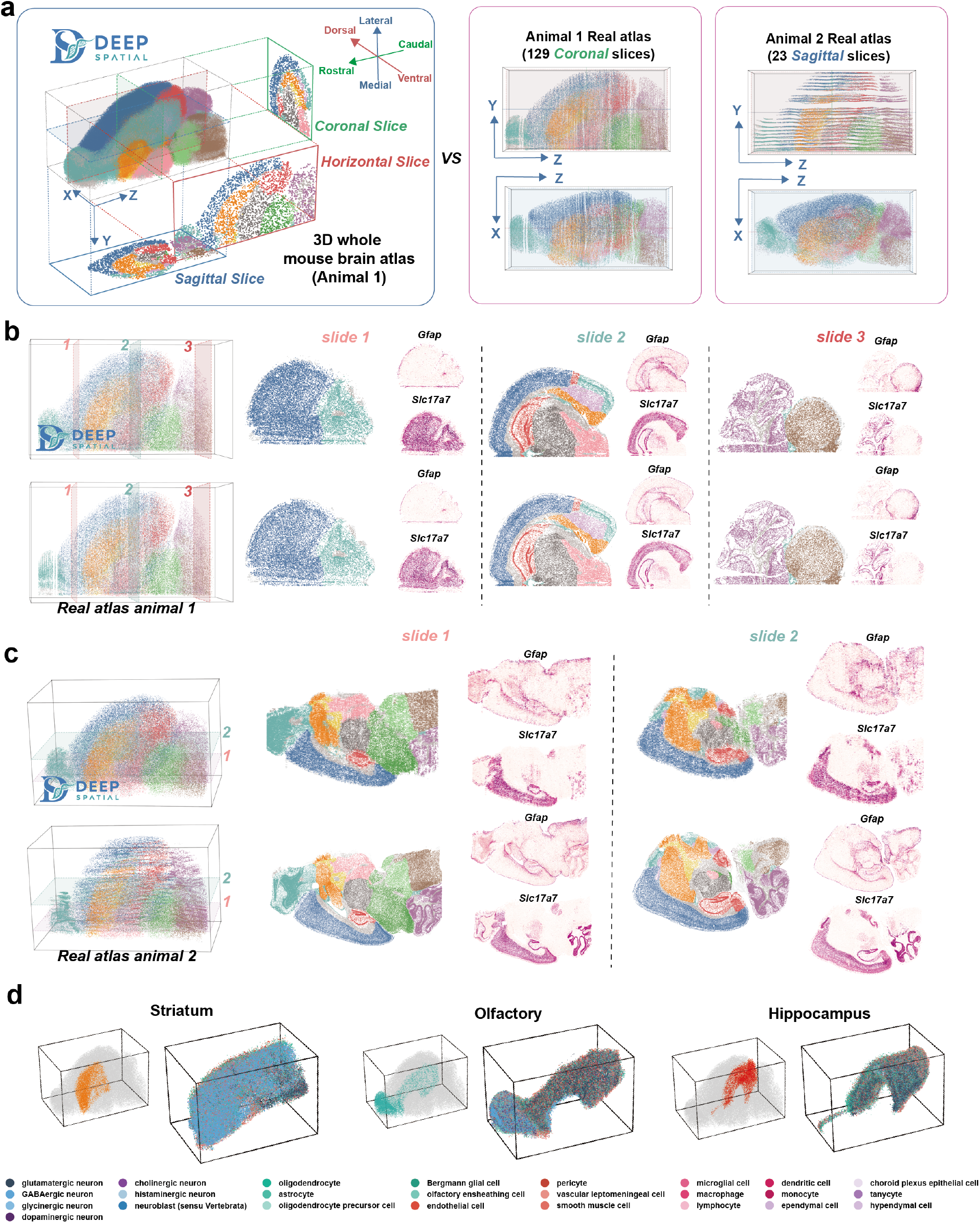
Multi-planar virtual slicing and 3D visualization of mouse whole-brain atlas. **a**, Schematic of 3D reconstruction and multi-planar validation. **b**, Coronal virtual slices versus ground-truth coronal sections. **c**, Horizontal virtual slices versus ground-truth horizontal sections. **d**, 3D visualization of cellular architecture in brain regions.

Notably, the reconstruction maintained high biological fidelity even in the challenging sagittal and horizontal planes. For horizontal slicing, we compared virtual slices from the reconstructed volume with both the genuine horizontal sections (Animal 2) and the horizontally stacked raw 129 coronal sections (Figs. 6c, S15a, b). The raw sparse coronal slices exhibited discontinuous, incomplete boundaries across large gaps in the horizontal direction, with severe structural fragmentation. In contrast, the DeepSpatial-reconstructed 3D volume reproduced continuous, anatomically consistent horizontal sections highly similar to genuine experimental horizontal slices, both in brain region organization and spatial gene expression patterns (Fig. 6c). Comparable results were observed in sagittal virtual slicing (Figs. S16a, b).

Beyond virtual sectioning, 3D reconstruction enables direct visualization of fine spatial architectures within individual brain regions. In the striatum, olfactory bulb, and hippocampus, we observed continuous volumetric organization and clear 3D spatial arrangements of fine cellular subtypes (Fig. 6d). Collectively, these results demonstrate that DeepSpatial captures realistic, fine-grained 3D cellular architecture compatible with multi-directional virtual slicing and direct volumetric analysis, supporting its generalizability for atlas-scale 3D reconstruction.

## 3 Discussion

The spatial omics community has established a highly mature ecosystem^35^ of computational tools dedicated to resolving planar heterogeneity, batch correction^36^, and spatial clustering^37^. Moving beyond these critical yet plane-restricted achievements, DeepSpatial is designed to tackle a completely different frontier: bridging the physical gaps inherently left by tissue sectioning. Our core contribution is fundamentally shifting the analytical paradigm from discrete 2.5D stacking to true topological continuity. This conceptual leap allows the seamless transformation of physically fragmented slices into an infinitely resolvable 3D atlas. Consequently, DeepSpatial functions as an essential infrastructural layer; it does not compete with current spatial pipelines, but rather empowers them by providing the continuous 3D foundation necessary for authentic whole-organ analysis.

The methodological superiority of DeepSpatial lies in its departure from rigid spatial registration and discrete linear interpolation. Unlike alignment-centric methods that merely correct geometric distortions, or imputation frameworks that insert pseudo-2D planes, DeepSpatial learns the underlying continuous dynamics of the tissue microenvironment. Our utilization of UOT establishes biologically rigorous cross-slice correspondences that naturally accommodate variations in tissue cross-sectional areas and cellular densities along the z-axis, relaxing the restrictive assumption of strict mass conservation. Coupled with a scalable multi-modal Transformer, the learned continuous vector field ensures that intermediate tissue states generated via ODE integration rigorously preserve both macroscopic cellular density and microscopic molecular gradients, completely eliminating the z-axis discontinuities inherent to previous 2.5D approaches.

A key strength of DeepSpatial lies in its true 3D reconstruction, which preserves biological continuity with high fidelity. This is evidenced by its superior spatial concordance with ground-truth 3D data compared to conventional 2.5D stacking approaches. Such fidelity enables the recovery of structures that are often missed due to sparse sectioning, most notably continuous vascular networks within tumors. In breast cancer IMC datasets, DeepSpatial reconstructs intact vasculature from fragmented sections, revealing spatial patterns such as the preferential localization of immune cells at distal vascular termini^38^. DeepSpatial is broadly compatible with both spatial proteomics (IMC) and transcriptomics platforms (including Deep-STARmap and openST), and remains robust under challenging conditions, such as irregular inter-slice spacing in HNSCC openST data, while preserving cell identity and marker expression profiles. Importantly, it also addresses key computational bottlenecks, as detailed in our comprehensive cost analysis (Table S3). For instance, the complete reconstruction of a 39-million-cell mouse whole-brain atlas was achieved in approximately 41.6 hours on a mid-range server. In stark contrast, conventional baseline methods such as SpatialZ would require an estimated 801 hours for the same scale, underscoring the exceptional efficiency of our approach. Collectively, the combination of biological fidelity, cross-platform applicability, and unprecedented computational scalability positions DeepSpatial as a practical solution for large-scale three-dimensional modeling of tissue microenvironments.

Despite these advancements, two primary limitations warrant consideration. First, the fidelity of the continuous 3D reconstruction is intrinsically tied to the quality of the upstream 2D input data and the initial spatial alignment. DeepSpatial is susceptible to technical noise inherent in spatial omics, such as imperfect cell segmentation, high transcriptomic dropout rates, or alignment artifacts, which can propagate through the generative ODE trajectory. Second, the accuracy of flow matching-based generation fundamentally depends on the spatial interval between adjacent slices. While the framework seamlessly interpolates standard gaps, extreme physical distances or abrupt, non-continuous morphological mutations (e.g., at sharp organ boundaries) remain challenging, potentially leading to over-smoothing or the generation of spurious transitional states.

Beyond the scope of these immediate technical constraints, DeepSpatial fundamentally redefines what can be extracted from spatial omics data. By providing whole 3D virtual tissue, the framework serves as a versatile computational platform for the development of next-generation downstream 3D analytical tools. This catalyzes a critical shift from traditional planar analyses to true volumetric quantification, unlocking the ability to rigorously map 3D cell-cell communication networks and topologically complex spatial domains. Furthermore, by expanding the continuous vector field to incorporate the temporal dimension, future iterations could seamlessly transition from 3D to true 4D spatio-temporal modeling. Such advancements will provide unprecedented mechanistic insights into highly dynamic biological processes, including organogenesis, tissue repair, and longitudinal tumor evolution. Ultimately, DeepSpatial establishes a robust, continuous analytical infrastructure, propelling spatial omics into the era of comprehensive whole-organ mapping.

## 4 Methods

### 4.1 Problem formulation

The fundamental objective of continuous 3D spatial omics reconstruction is to generate a cohesive 3D tissue volume, denoted as *V*_01_, from two discrete, adjacent 2D slices, denoted as *S*_0_ and *S*_1_. These source slices are located at physical *Z*-axis coordinates *z*_0_ and *z*_1_, respectively, and can be represented as empirical distributions of cells. Within either slice, each cell *i* is characterized by a state tuple *s*_*i*_ = (**x**_*i*_, **g**_*i*_, *c*_*i*_), where **x**_*i*_ ∈ ℝ^2^ denotes the 2D spatial coordinates, **g**_*i*_ ∈ ℝ^*D*^ is the high-dimensional gene expression vector (*D* being the number of genes), and *c*_*i*_ ∈ {1,…, *K*} represents the discrete cell type label. Crucially, to construct the continuous volume *V*_01_, every generated virtual cell must be assigned a strict 3D spatial coordinate (**x**, *z*) ∈ ℝ^3^, alongside its corresponding inferred gene expression **g** and cell type *c*.

We formulate this *S*_0_, *S*_1_ → *V*_01_ interpolation mathematically as learning a continuous vector field via flow matching. The specific implementation is decoupled into two phases. First, we learn a continuous, time-dependent vector field **v**_*t*_ that models the underlying dynamics transporting the joint distribution of cell states *s* = (**x, g**, *c*) from the source slice *S*_0_ at *z*_0_ to the target slice *S*_1_ at *z*_1_. Second, to populate the continuous volume *V*_01_, we sample from this learned vector field. For any arbitrary intermediate physical coordinate *z*_*t*_ ∈ (*z*_0_, *z*_1_), we sample the virtual cells and infer their multidimensional states through ODE integration, effectively assembling the discrete 2D inputs into a highly resolved 3D microenvironment.

### 4.2 Data alignment and setup

To construct a coherent 3D microenvironment from sparsely sampled 2D sections, it is imperative to first align the discrete source slices into a common spatial coordinate system. For datasets equipped with a pre-registered coordinate framework (e.g., the Allen Mouse Brain CCFv3^39^), we directly adopt the registered spatial coordinates. Conversely, for datasets lacking a pre-registered system, we apply pairwise spatial alignment algorithms, such as PASTE^16^, which integrate high-dimensional gene expression similarity and physical spatial proximity to robustly align cells across adjacent slices.

Following this initial spatial alignment, we set up the computational domain to ensure numerical stability during neural network training and ODE integration. Specifically, we project the aligned physical coordinates into a standardized, dimensionless normalization space. The aligned 2D spatial coordinates **x** ∈ ℝ^2^ and the physical depth *z* across all input slices (e.g., *S*_0_ and *S*_1_) are globally normalized to the range [0, 1] using a min-max scaling approach. We record the global extreme values and corresponding physical spans for the entire measured tissue volume to perform this linear transformation. This system setup guarantees that the continuous vector field **v**_*t*_ is learned within a well-conditioned, isotropic coordinate system, strictly mitigating artifacts induced by arbitrary physical measurement scales. Upon generating the continuous volume *V*_01_, the physical space is exactingly restored using the inverse transformation, mapping the normalized generated coordinates back to their absolute physical dimensions to assign precise, metric-accurate 3D coordinates to all virtual cells.

### 4.3 Establishing Optimal Transport for Cell Coupling

To construct the target probability paths for flow matching, we must first establish biologically meaningful cell-to-cell correspondences between adjacent slices^40^. We represent the cell distributions of the preprocessed source and target slices, *S*_0_ and *S*_1_, as empirical probability measures *µ*_0_ and *µ*_1_. To account for variations in cell density, tissue morphology, and the biological realities of cell proliferation or apoptosis between slices, we formulate this alignment as an UOT^26^.

Let *N*_0_ = |*S*_0_| and *N*_1_ = |*S*_1_| denote the number of cells in the respective slices. To guide the optimal transport, we construct a comprehensive cost matrix 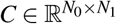 that rigorously integrates spatial proximity, transcriptomic similarity, and categorical cell-type consistency. Specifically, we compute the normalized Euclidean spatial distance matrix *D*^spat^ and the normalized Cosine gene expression distance matrix *D*^gene^, where each distance matrix is divided by its maximum value to ensure scale invariance. Furthermore, to biologically restrict matching between disparate cell lineages, we introduce a cell-type penalty matrix 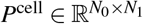, defined using the indicator function as 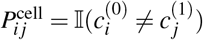. The total cost matrix is thus formulated as:

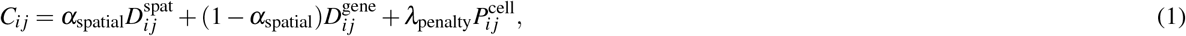

where *α*_spatial_ ∈ [0, 1] balances the spatial and transcriptomic modalities, and *λ*_penalty_ is assigned to strictly penalize cross-cell-type couplings.

The soft coupling matrix 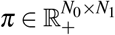 is subsequently obtained by minimizing the entropy-regularized UOT objective:

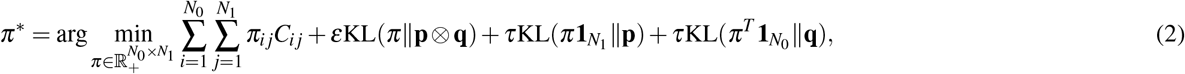

where **p** and **q** represent the uniform marginal distributions of *S*_0_ and *S*_1_. The parameter *ε* controls the entropic regularization, ensuring computational tractability and differentiability via the Sinkhorn algorithm^41,42^, while *τ* dictates the marginal relaxation, mathematically penalizing mass creation and destruction, which corresponds biologically to cellular proliferation and apoptosis within the tissue.

### 4.4 Neural Architecture and Continuous flow matching Formulation

To effectively model the complex spatio-transcriptomic dynamics over the continuous physical depth, we parameterize the time-dependent vector field using the Gene Diffusion Transformer (GiT), a scalable, multi-modal Transformer architecture^27,28,43^. The core of GiT lies in its unified tokenization strategy, which translates heterogeneous physical and biological modalities (namely spatial coordinates, gene expressions, and cell types) into a shared, high-dimensional representation space.

Given a continuous integration time *t* ∈ [0, 1], we decouple the multidimensional cell state into distinct token streams. The 2D spatial coordinate **x**_*t*_ ∈ ℝ^2^ is linearly projected into a single spatial token **h**^(*x*)^ ∈ ℝ^1×*H*^, where *H* is the hidden dimension. Concurrently, the high-dimensional transcriptomic vector **g**_*t*_ ∈ ℝ^*D*^ is chunked into discrete patches and processed by a non-linear Multi-Layer Perceptron (MLP) embedder, yielding a sequence of gene tokens **H**^(*g*)^ ∈ ℝ^*M*×*H*^, where *M* = ⌈*D/*patch_size⌉. These token streams are concatenated and augmented with a 1D sinusoidal positional embedding **E**_pos_ to form the unified input sequence **H** = [**h**^(*x*)^; **H**^(*g*)^] + **E**_pos_ ∈ ℝ^(1+*M*)×*H*^.

Instead of appending the categorical cell type directly to the sequence, GiT encodes the one-hot cell lineage anchor **c**_*t*_, alongside the temporal state *t* and physical depth constraints (*z*_*t*_, Δ*z*), into a unified global context token **w**_*t*_ ∈ ℝ^*H*^ :

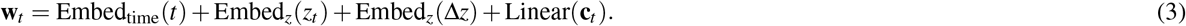

This global token deeply modulates the sequence **H** through a backbone of Transformer blocks equipped with adaptive Layer Normalization (adaLN-Zero). For each block, the global context **w**_*t*_ dynamically regresses the scale (*γ*), shift (*β*), and gate (*α*) parameters, dictating how the spatial and gene tokens interact via Multi-Head Self-Attention (MSA) and MLP sub-layers:

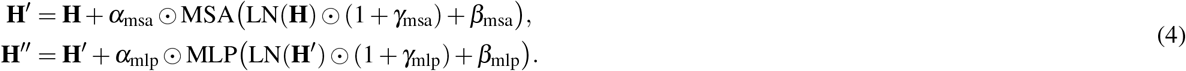

All modulation layers are strictly zero-initialized, guaranteeing that the network functions as an identity mapping at the onset of training to stabilize the underlying Ordinary Differential Equation (ODE) integration^29^.

Through task-specific output heads, the processed sequence is projected back to their respective physical and biological domains. Crucially, we formulate the generative process as a mixed continuous-discrete flow matching framework^25^ to accommodate the multi-modal nature of the cell state. Under the flow matching framework, we define a deterministic linear probability path for the continuous spatial and transcriptomic modalities, yielding constant target velocities (i.e., **x**_1_ − **x**_0_ and **g**_1_ − **g**_0_), while simultaneously modeling the categorical cell type as a discrete probability flow. The unified objective function jointly optimizes these heterogeneous modalities by minimizing the expected vector field matching error for continuous states and the cross-entropy for discrete states:

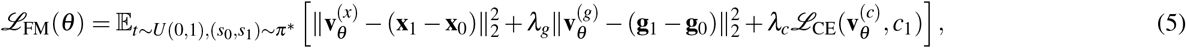

where the spatial velocity 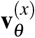 is decoded from the first sequence token, the transcriptomic velocity 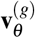 is decoded from the remaining *M* tokens, and the cell-type evolution 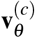 is derived from the mean-pooled representation of the entire sequence. The coefficients *λ*_*g*_ and *λ*_*c*_ rigidly balance the continuous regression and discrete classification tasks within the shared generative vector field.

### 4.5 Generating the 3D Volume via ODE Integration

To generate the intermediate microenvironment at an arbitrary physical depth *z*_*t*_ ∈ (*z*_0_, *z*_1_), we construct the generative trajectory by solving the ODE defined by our continuous vector field. To minimize numerical integration error and equilibrate boundary information from adjacent slices, we implement a bidirectional sampling strategy. For each synthesized cell, we introduce a Bernoulli random variable *b* ∼ Bernoulli(0.5) to determine the direction of evolution. The generated continuous state *s*_*t*_ = (**x**_*t*_, **g**_*t*_) is obtained via the following piecewise Lebesgue integration:

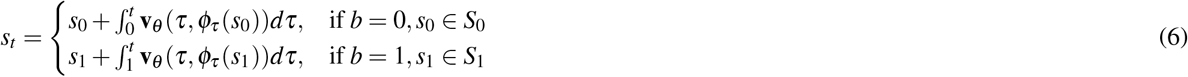

where *t* ∈ (0, 1) is the fractional integration time corresponding to *z*_*t*_, and *φ*_*τ*_ denotes the flow map. This initial value problem is numerically approximated using an ODE solver (e.g., the Dormand-Prince method^44^) to guarantee local error bounds and computational efficiency.

To computationally render a 3D tissue volume that rigorously preserves the macroscopic cellular density gradients, we apply inverse transform sampling over the continuous temporal measure. Assuming the cellular density rate *ρ*(*τ*) interpolates linearly such that *ρ*(*τ*) = (1 − *τ*)*N*_0_ + *τN*_1_, we formulate the cumulative distribution function (CDF) *F*(*t*) as the normalized integral of the density measure:

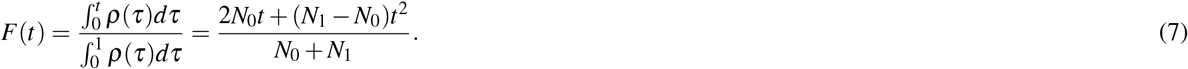

By equating this CDF to a uniform random variable *u* ∼ *U* (0, 1) and isolating *t*, we derive the exact fractional temporal position *t* = *F*^−1^(*u*) via the positive quadratic root:

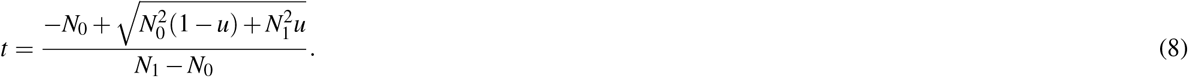

Following the ODE integration, which yields an oversampled raw virtual volume 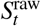, we apply a spatial pruning optimization to eliminate out-of-distribution artifacts and enforce structural fidelity. The pruned set 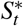 is extracted by solving a subset minimization problem that retains exactly *N*(*t*) cells exhibiting the minimum expected spatial transport cost to their respective source anchors:

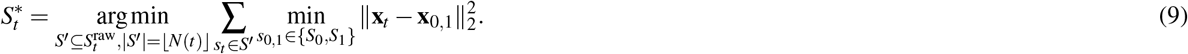

This optimization guarantees that the synthesized 3D volume remains spatially consistent with the empirical tissue morphology while strictly adhering to the estimated cellular abundance at each depth.

### 4.6 Downstream Analysis of Continuous 3D Microenvironments

To construct virtual slices from the 3D spatial data, we projected the cells within a 10 µm range along the Z-axis onto a 2D plane. Specifically, for each point on the Z-axis, we selected a 5 µm thick slice of cells both above and below the given point and performed a 2D projection onto the corresponding screen to make a virtual slice.

We compared the reconstructed 3D spatial tissue with the real 3D dataset by first calculating voxel compositions for both. Voxel compositions were obtained by normalizing the spatial coordinates and mapping them to a 3D grid. Each voxel contained the relative proportion of cell types, computed using one-hot encoding for the predicted labels. To quantify the similarity between the reconstructed and real datasets, we computed the cosine similarity between their voxel compositions:

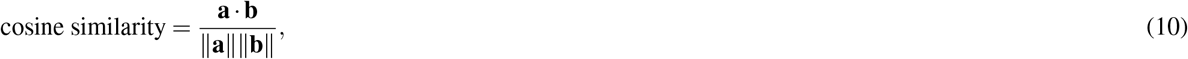

where **a** and **b** are the normalized voxel composition vectors for the ground truth and predicted datasets, respectively^45^. The resulting similarity map was visualized in 3D, allowing for a direct comparison of the accuracy of the reconstruction.

In gene-level comparative analysis, Moran’s *I* statistic was employed to quantify the spatial autocorrelation of gene expression, defined as:

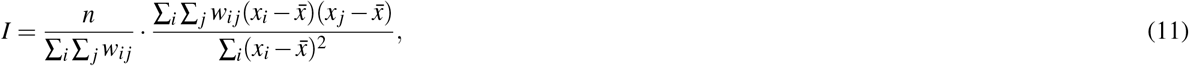

where *n* denotes the number of observations, *w_ij_* is the spatial weight matrix, *x*_*i*_ represents the gene expression value, and 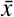 is the mean expression.

We selected the top 100 highly variable genes from the original dataset and performed Spearman correlation-based regression analyses between the original and reconstructed datasets to assess the preservation of gene expression patterns.

### 4.7 Downstream Analysis of BRCA IMC dataset

We first annotated each tissue section based on protein expression levels. Single-cell marker expression was range-normalized to the 99th percentile per channel across all cells, followed by clustering using the PhenoGraph with Manhattan distance and 10 nearest neighbors^46^. Cells were subsequently annotated into distinct cell types, including B cells, T cells, plasma cells, myeloid cells, fibroblasts, endothelial cells, basal epithelial cells, and luminal epithelial cells; unclassified cells were labeled as unknown.

Spatial neighborhood analysis of cell types was subsequently performed using Squidpy, where a graph network was built based on the 30 *µ*m neighboring cells^47^. We focused on cell populations adjacent to endothelial cells, and quantified the density of cell populations within a 30-*µ*m radius of endothelial cells at 5-*µ*m intervals along the z-axis.

### 4.8 Downstream Analysis of openST HNSCC metastatic lymph node datasets

For the full-transcriptome HNSCC spatial dataset, we selected the top 2000 highly variable genes and performed 3D reconstruction. A common overlapping region was defined via the maximum coincident rectangle (X-range: [4991.99, 11238.94]; Y-range: [371.53, 8071.74]) to account for variable slice ranges. The cellular proportion distribution was calculated at 5-*µ*m intervals along the z-axis, representing the fractional composition of each cell subtype and spatial domain within each layer. For comparison, the cellular proportions in the corresponding actual tissue sections were computed within individual slices. Cellular neighborhood analysis in three-dimensional (3D) tissues was defined by computing the local cellular density within a spherical region of radius *r* = 20 *µ*m centered on each tumor cell:

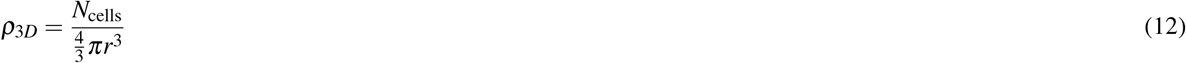

where *N*_cells_ denotes the number of neighboring cells within the sphere.

For two-dimensional (2D) sections with a thickness of *h* = 10 *µ*m, the neighborhood was approximated by a cylindrical volume with base radius *r* = 20 *µ*m and height equal to the section thickness:

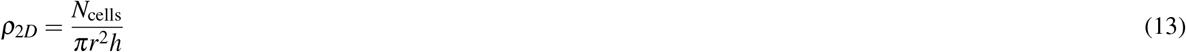

where *N*_cells_ represents the number of cells within the cylindrical region.

Gene embeddings were extracted using scVI, spatial domains were identified with CellCharter, and a 3D spatial atlas was constructed^34^.

### 4.9 Downstream Analysis of BRAIN Initiative Cell Census Network data

After being aligned to the Allen Mouse Brain CCFv3, the BRAIN Initiative Cell Census Network data were rendered as multi-slice tissues with a 100-*µ*m interval. These slices were subsequently reconstructed into 3D tissues via the DeepSpatial pipeline. The multidirectional distribution of cell type proportions represents the spatial arrangement of cells within the layer at a 5-*µ*m interval along the corresponding axis. The reconstructed 3D tissue was randomly downsampled according to the proportion of each brain region to match the total cell count (4,167,869) of the original real multi-slice dataset. We then employed SCVI for gene expression feature extraction and UMAP dimensionality reduction^36^, to evaluate the correspondence of cell types and brain regions between the reconstructed tissue and the original slices. For the comparison of local microenvironments, we quantified the cellular compositional proportion of each major brain region and calculated the Spearman correlation between the 3D reconstructed tissue and the real multi-slice data. We also performed multidirectional virtual slice comparisons, where the virtual slices were set to a thickness of 10 *µ*m to maintain consistency with the real slices.

The comprehensive hyperparameter configurations for the entire DeepSpatial pipeline, including optimal transport, neural network architecture, flow matching, and ODE integration, are detailed in Table S1.

## Author contributions statement

Conceptualization: Y.H.Y., Y.M.L., K.Z., Y.C., L.S. and E.H.C.; Methodology: Y.H.Y. and Y.M.L.; Software: Y.H.Y. and Y.G.B.; Formal Analysis: Y.H.Y. and Y.M.L.; Investigation: Y.H.Y., Y.M.L. and K.Z.; Data Curation: Y.M.L.; Validation: Y.H.Y. and Y.M.L.; Visualization: Y.H.Y. and Y.M.L.; Writing – Original Draft: Y.H.Y. and Y.M.L.; Writing – Review & Editing: K.Z., Z.X., H.X.P., R.Y., Q.L., D.F.L., Y.C., L.S. and E.H.C.; Supervision: K.Z., Y.C., L.S. and E.H.C.; Project Administration: K.Z.; Resources: K.Z., Y.C., L.S. and E.H.C.; Funding Acquisition: K.Z. and E.H.C. All authors reviewed the manuscript.

## Data availability

The spatial omics datasets analyzed in this study comprise Deep-STARmap (human cSCC, mouse brain), Deep-RIBOmap (mouse brain)^13^, breast cancer imaging mass cytometry (IMC)^11^, openST HNSCC metastatic lymph node^33^, and BRAIN Initiative mouse brain MERFISH datasets^10^, all acquired from public spatial omics repositories. Detailed information on dataset platforms and sample statistics is summarized in Table S2.

All processed data generated during this study are available from the corresponding author on a reasonable request.

## Code availability

The project overview and interactive documentation of DeepSpatial are accessible at https://yyh030806.github.io/DeepSpatial. The full source code, model implementation, training/inference pipelines, and reproducibility tutorials are openly available under an open-source license at https://github.com/yyh030806/DeepSpatial.

## Notes

### Competing Interest Statement

The authors have declared no competing interest.

### Summary of Updates

Graphical abstract added to summarize main findings; author list and affiliations updated to include new contributors.

