## Supplementary Information for "Reconstructing True 3D Spatial Omics at Single-Cell Resolution"

#### Supplementary contents

##### Supplementary Notes

- Supplementary Note S1 – Overview of the related 2.5D baseline methods.
- Supplementary Note S2 – Mathematical derivations of the DEEPSPATIAL algorithm.
- Supplementary Note S3 – Detailed architecture of the Gene Diffusion Transformer (GiT).

##### Supplementary Tables

- Supplementary Table S1 – Hyperparameter configurations of DEEPSPATIAL.
- Supplementary Table S2 – Summary of the evaluated datasets.
- Supplementary Table S3 – Computational cost analysis of DEEPSPATIAL.

##### Supplementary Figures

- Supplementary Figure S1 – Detailed spatial and transcriptomic characteristics of the 3D human cSCC Deep-STARmap dataset.
- Supplementary Figure S2 – Detailed spatial and transcriptomic characteristics of the 3D mouse brain Deep-STARmap dataset.
- Supplementary Figure S3 – Detailed spatial and transcriptomic characteristics of the 3D mouse brain Deep-RIBOmap dataset.
- Supplementary Figure S4 – Spatial distribution of cell populations across all experimental slices in the breast cancer (BRCA) IMC dataset.
- Supplementary Figure S5 – Proportions of cell-type composition across all experimental slices in the BRCA IMC dataset.
- Supplementary Figure S6 – Gene expression heatmaps across all experimental slices in the BRCA IMC dataset.
- Supplementary Figure S7 – Comprehensive comparison of vasculature-associated features between DEEPSPATIAL-reconstructed synthetic tissue and experimental BRCA IMC slices.
- Supplementary Figure S8 – Vertical virtual slice comparisons between DEEPSPATIAL-reconstructed synthetic tissue and experimental BRCA IMC data.
- Supplementary Figure S9 – Cellular spatial architecture of the openST HNSCC metastatic lymph node multi-slice dataset.
- Supplementary Figure S10 – Gene expression profiles of distinct cell types between DEEPSPATIAL-reconstructed synthetic data and original openST HNSCC dataset.
- Supplementary Figure S11 – Comparison of cell-type distribution within spatial domains between DEEPSPATIAL-reconstructed synthetic data and original openST HNSCC dataset.
- Supplementary Figure S12 – 3D spatial domain architecture of the DEEPSPATIAL-reconstructed openST HNSCC dataset.
- Supplementary Figure S13 – Gene expression profiles of distinct cell types between DEEPSPATIAL-reconstructed synthetic data and the BRAIN Initiative Cell Census Network mouse brain dataset.
- Supplementary Figure S14 – Cell-type and brain region distribution along the X/Y axes between DEEPSPATIAL-reconstructed synthetic data and the BRAIN Initiative Cell Census Network mouse brain dataset.
- Supplementary Figure S15 – Horizontal virtual slice comparison of coronal planes between DEEPSPATIAL-reconstructed synthetic data and the BRAIN Initiative Cell Census Network real mouse brain data.
- Supplementary Figure S16 – Sagittal virtual slice comparison of coronal planes between DEEPSPATIAL-reconstructed synthetic data and the BRAIN Initiative Cell Census Network real mouse brain data.

### Supplementary Notes

#### S1 Overview of Related 2.5D Baseline Methods

Current computational frameworks for reconstructing three-dimension (3D) spatial omics from 2D multi-slice datasets can be broadly categorized into alignment-based methods and generative-based imputation methods.

**Alignment-Based Methods**. Alignment-based methods primarily focus on projecting discrete, consecutive 2D slices into a common spatial coordinate system using rigid or non-rigid transformations. The representative baseline methods include:

- **PASTE**[16]: Utilizes Fused Gromov-Wasserstein Optimal Transport to align adjacent slices by simultaneously considering physical spatial distances and high-dimensional transcriptional similarities.
- **GPSA**[17]: Employs deep Gaussian processes to map multi-layered two-dimensional spatial omics sections into a unified, predictable 3D coordinate system.
- **CAST**[14]: Leverages graph convolutional networks to extract spatially-aware transcriptomic features, enabling robust cross-scale alignment and integration of tissue sections.
- **ELD**[15]: Uses deep learning frameworks for spatial landmark detection, facilitating non-rigid tissue registration and structural alignment across consecutive slices.

While these methods successfully establish spatial correspondence across measured sections, they inherently function as 2.5D stacking approaches. They solely manipulate existing data planes without inferring the missing biological information within the physical gaps between slices. Consequently, the resulting representations suffer from dimensional fragmentation, failing to recover the continuous tissue topology and fine, depth-dependent molecular gradients that the DEEPSPATIAL successfully resolves.

**Generative-Based Methods** To overcome the sparse sampling limits of physical sectioning, recent generative methods have advanced beyond simple alignment to active data imputation, inferring virtual representations to fill the physical gaps. The representative baselines include:

- **SpatialZ**[18]: A pioneering framework that imputes intermediate virtual slices between adjacent experimental sections, attempting to bridge the planar-to-3D gap by assembling a denser 2.5D tissue stack.
- **MIMYR**[19]: Utilizes generative modeling techniques specifically designed to synthesize and impute structurally missing tissue regions across incomplete spatial omics datasets.
- **Morphe**[20]: Bridges the modalities of image generation and spatial omics to synthesize spatially resolved virtual tissue representations guided by image-conditioned generative models.

Despite significant advancements in virtual imputation, these generative frameworks remain fundamentally constrained to the 2.5D paradigm. Methods like SpatialZ construct denser volumes by inserting discrete planar slices, yet the underlying representation remains a fragmented stack. Even with dense virtual interpolation, this layer-by-layer insertion forcibly disrupts the continuous spatial connectivity along the Z-axis. Such discrete frameworks fundamentally fail to capture the fluid, uninterrupted topological transitions within biological microenvironments, which strictly necessitates the infinitely resolvable, true 3D continuous ODE modeling achieved by DEEPSPATIAL.

#### S2 Theoretical Derivations of the DEEPSPATIAL Algorithm

This section provides the intermediate mathematical proofs and algorithmic solving steps that are omitted in the main text Methods section due to length constraints.

**Sinkhorn Iterations for Unbalanced Optimal Transport.** In the main text (Eq. 2), we formulated the entropy-regularized Unbalanced Optimal Transport (UOT) objective. Due to the strictly convex nature of this entropy-regularized objective, the optimal soft coupling matrix $\pi^*$ cannot be solved analytically but is efficiently computed using the generalized Sinkhorn-Knopp algorithm.

By introducing the Gibbs kernel matrix $K = \exp(-C/\varepsilon)$, where $C$ is the total cost matrix (Eq. 1), the optimal coupling can be factorized as $\pi^* = \text{diag}(u)K\text{diag}(v)$. The vectors $u \in \mathbb{R}^{N_0}$ and $v \in \mathbb{R}^{N_1}$ are dual scaling vectors, which are iteratively updated via:

$$u^{(iter+1)} = \left(\frac{p}{Kv^{(iter)}}\right)^{\frac{\tau}{\tau+\varepsilon}}, \quad v^{(iter+1)} = \left(\frac{q}{K^T u^{(iter+1)}}\right)^{\frac{\tau}{\tau+\varepsilon}}$$

where $p$ and $q$ represent the uniform marginal distributions of the source and target slices. Convergence of these alternating updates yields the optimal sparse coupling matrix $\pi^*$, which effectively provides the biologically rigorous cell-to-cell correspondences required for sampling the downstream generative trajectories.

**Integral Derivation for Density-Preserving Sampling.** In the main text (Eq. 7 and 8), we present the closed-form solutions for the Cumulative Distribution Function $F(t)$ and the fractional temporal position $t$. Here we provide the detailed piecewise integration steps.

Assuming the cellular density rate $\rho(\tau)$ interpolates linearly between the two measured slices: $\rho(\tau) = (1-\tau)N_0 + \tau N_1$. The denominator, representing the total expected cells across the continuous volume, is integrated as:

$$\int_0^1 \rho(\tau)d\tau = \left[N_0\tau - \frac{1}{2}N_0\tau^2 + \frac{1}{2}N_1\tau^2\right]_0^1 = \frac{N_0+N_1}{2} \tag{14}$$

The numerator, representing the cumulative density up to a given fractional time $t \in [0,1]$, is calculated as:

$$\int_0^t \rho(\tau)d\tau = \left[N_0\tau - \frac{1}{2}N_0\tau^2 + \frac{1}{2}N_1\tau^2\right]_0^t = N_0 t + \frac{1}{2}(N_1 - N_0)t^2 \tag{15}$$

Dividing the numerator by the denominator directly yields the exact cumulative distribution function (CDF) formulation $F(t)$ shown in Eq. 7.

To perform inverse transform sampling using a uniform random variable $u \sim U(0,1)$, we equate $F(t) = u$. Multiplying both sides by the denominator $(N_0 + N_1)$ and rearranging the terms yields the standard quadratic equation with respect to $t$:

$$(N_1 - N_0)t^2 + 2N_0 t - (N_0 + N_1)u = 0 \tag{16}$$

Applying the quadratic formula and isolating the valid positive root $t \in [0,1]$ directly results in the exact temporal sampling position $t$ presented in Eq. 8 of the main text.

#### S3 Detailed Architecture of Gene Diffusion Transformer (GiT)

While the main text formulates the mathematical forward pass of the Gene Diffusion Transformer (GiT), this section provides the concrete architectural and engineering details necessary for implementation and reproducibility. The GiT architecture is inspired by Diffusion Transformers but heavily customized to process heterogeneous multi-modal spatial omics data within a unified continuous-discrete flow matching framework.

**Tokenization and Feature Embeddings.** The foundation of GiT is its unified tokenization strategy, which projects disparate modalities into a shared hidden dimension $H$.

- **Spatial and Transcriptomic Tokenization:** The 2D spatial coordinate $\mathbf{x}_t$ is projected into a single token $\mathbf{h}^{(x)}$ via a multi-layer perceptron (MLP) consisting of two linear layers with a SiLU (Swish) activation. For the transcriptomic vector $\mathbf{g}_t \in \mathbb{R}^D$, we chunk the array into patches. These patches are processed by a non-linear embedder, specifically a 2-layer MLP with SiLU activation, yielding $M$ gene tokens.
- **Positional Encoding:** To inject structural awareness, we utilize standard 1D fixed sinusoidal positional embeddings $\mathbf{E}_{pos}$. These embeddings are added directly to the concatenated sequence of the spatial and gene tokens before entering the Transformer blocks.
- **Global Context Embedding:** As defined in Eq. 3 of the main text, the global context token $\mathbf{w}_t$ conditions the network. The continuous temporal state $t \in [0,1]$ and physical depth constraints $(z_t, \Delta z)$ are first mapped into higher-dimensional representations using sinusoidal Fourier feature embeddings, followed by a linear projection. The categorical cell lineage anchor $\mathbf{c}_t$ is embedded via a standard lookup table (nn.Embedding). These four components are summed to form $\mathbf{w}_t$.

**adaLN-Zero Modulated Transformer Blocks.** The core processing backbone consists of stacked Transformer blocks equipped with adaptive Layer Normalization (adaLN-Zero). Unlike standard layer normalization, the scale ($\gamma$), shift ($\beta$), and gate ($\alpha$) parameters are not learnable scalar weights. Instead, they are dynamically regressed from the global context token $w_t$ using a conditioning MLP with SiLU activations.

Crucially, to ensure that the initial ODE integration trajectories remain well-conditioned, we employ the *zero-initialization* trick. The final linear layer of the conditioning MLP that outputs the $\alpha$, $\gamma$, and $\beta$ parameters is strictly initialized to zero. This guarantees that at the start of training, the Transformer blocks function as exact identity mappings, significantly stabilizing the early phases of flow matching optimization.

**Task-Specific Output Decoders.** After processing through the sequence of Transformer blocks, the output sequence must be decoupled and mapped back to the physical and biological target spaces to generate the velocity vectors.

- **Continuous Spatial Decoder:** The 0-th token of the output sequence, corresponding to the spatial position, is passed through a linear projection head to predict the 2D spatial velocity $\mathbf{v}_\theta^{(x)}$.
- **Continuous Transcriptomic Decoder:** The subsequent $M$ tokens are passed through a shared linear projection head. The outputs are then reshaped and flattened to reconstruct the $D$-dimensional transcriptomic velocity vector $\mathbf{v}_\theta^{(g)}$.
- **Discrete Cell-Type Decoder:** Rather than relying on a single token, we apply average pooling across the entire processed sequence (spatial + gene tokens) to capture global microenvironmental features. This pooled representation is passed through a classification head (a linear layer) to output the unnormalized logits $\mathbf{v}_\theta^{(c)}$ over the $K$ cell categories, guiding the discrete probability flow.

#### Supplementary Tables

| Component | Hyperparameter | Description | Value |
|---|---|---|---|
| UOT Alignment | $\alpha_{spatial}$ | Weight for spatial vs. transcriptomic distance | 0.5 |
| | $\lambda_{penalty}$ | Penalty weight for cross-cell-type matching | 10.0 |
| | $\varepsilon$ (uot_reg) | Entropic regularization parameter for Sinkhorn | 0.8 |
| | $\tau$ (uot_tau) | Marginal relaxation parameter | 0.05 |
| | n_samples_base | Base number of cell pairs sampled per slice pair | Cell nums |
| Network (GiT) | patch_size | Tokenization patch size for spatial coordinates | 8 |
| | hidden_size | Transformer embedding dimension | 256 |
| | depth | Number of transformer layers | 6 |
| | num_heads | Number of attention heads | 8 |
| | mlp_ratio | Expansion ratio for MLP layers | 4.0 |
| flow matching | path_type | Probability path formulation | Linear |
| | $\lambda_g$ | Weight for continuous gene expression regression | 0.1 |
| | $\lambda_c$ | Weight for discrete cell type classification | 10.0 |
| | train_eps | Temporal boundary padding during training | 0.02 |
| | ema_decay | Exponential moving average (EMA) decay rate | 0.999 |
| ODE Integration | Solver (sampling_method) | Numerical integration method | dopri5 |
| | atol | Absolute tolerance for the ODE solver | 1e-5 |
| | rtol | Relative tolerance for the ODE solver | 1e-5 |
| | sample_eps | Temporal boundary padding during inference | 0.02 |
| | steps | Integration steps | 100 |
| Optimization | Optimizer | Network weight updating algorithm | AdamW |
| | lr | Initial learning rate | 2e-4 |
| | epoch | Number of epochs for training | 10 |
| | weight_decay | L2 regularization term for the optimizer | 1e-5 |
| | num_workers | Multi-process data loading workers | 4 |

**Table S1. Hyperparameter Configurations.** Detailed settings for data sampling, the Gene Diffusion Transformer (GiT) architecture, continuous flow matching objective, ODE integration heuristics, and network optimization utilized in the DEEPSPATIAL pipeline.

| Dataset Name | Tissue | Technology | Slices / Spacing / Thickness | Total Cells | Features |
|---|---|---|---|---|---|
| human cSCC | Squamous Cell Carcinoma | Deep-STARmap | - | 43,818 | 254 |
| mouse brain | Brain | Deep-STARmap | - | 198,675 | 1,017 |
| mouse brain | Brain | Deep-RIBOmap | - | 164,029 | 1,017 |
| BRCA IMC | Breast Cancer | IMC | 15 slices / 20 $\mu$m / 2 $\mu$m | 106,706 | 25 |
| openST HNSCC | Metastatic Lymph Node | openST | 19 slices / 30 $\mu$m / 10 $\mu$m | 1,097,769 | 2,000 |
| merfish brain | Brain | MERFISH | 129 slices / 100 $\mu$m / 10 $\mu$m | 4,167,869 | 1,120 |

**Table S2. Summary of Evaluated Datasets.** Overview of the spatial omics datasets used for 3D reconstruction and benchmarking.

| Dataset Name | Generated Cells | Training VRAM | Training Time | Inference VRAM | Inference Time |
|---|---|---|---|---|---|
| Human cSCC | 36,259 | 2.3 GB | 1.3 mins | 1.2 GB | 0.3 mins |
| Mouse brain | 202,052 | 8.3 GB | 18.8 mins | 4.4 GB | 11.6 mins |
| Mouse brain | 166,696 | 8.3 GB | 15.2 mins | 4.3 GB | 4.9 mins |
| BRCA IMC | 991,181 | 0.5 GB | 1.3 mins | 0.3 GB | 0.9 mins |
| openST HNSCC | 3,148,610 | 21.4 GB | 3.2 hours | 11.2 GB | 7.3 hours |
| Mouse brain | 39,168,380 | 23.3 GB | 16.0 hours | 13.8 GB | 25.6 hours |

**Table S3. Computational Cost.** Training and inference time and VRAM overhead across various datasets on an NVIDIA Tesla V100 GPU.

### Supplementary Figures

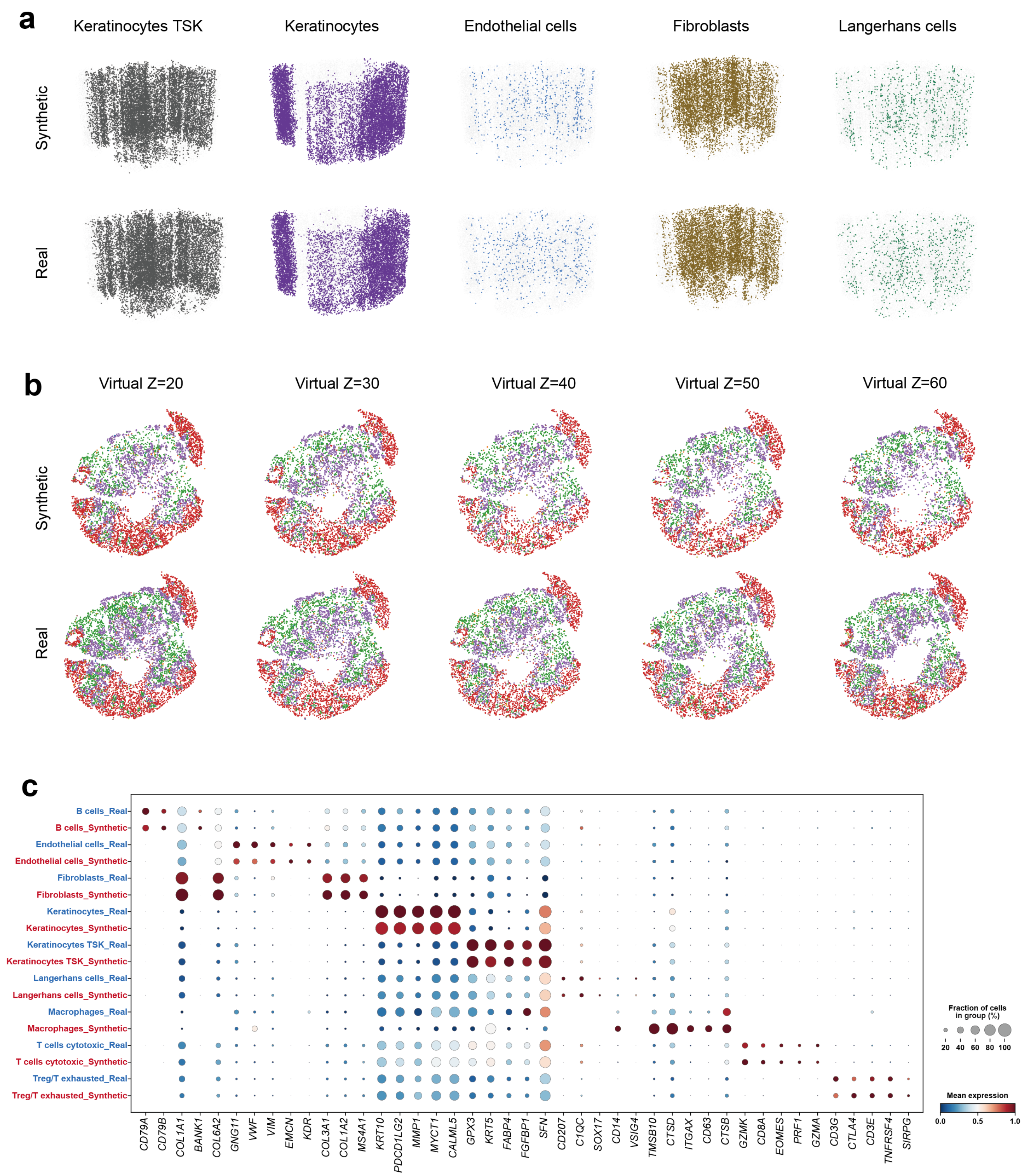


**Fig. S1.** Detailed spatial and transcriptomic characteristics of the 3D human cSCC Deep-STARmap dataset. **a,** 3D spatial distribution of distinct cell types in the cSCC dataset. **b,** Virtual tissue slices generated by DEEPSPATIAL (synthetic) versus ground-truth real 3D tissue slices. **c,** Dot plot displaying gene expression patterns across different cell types.

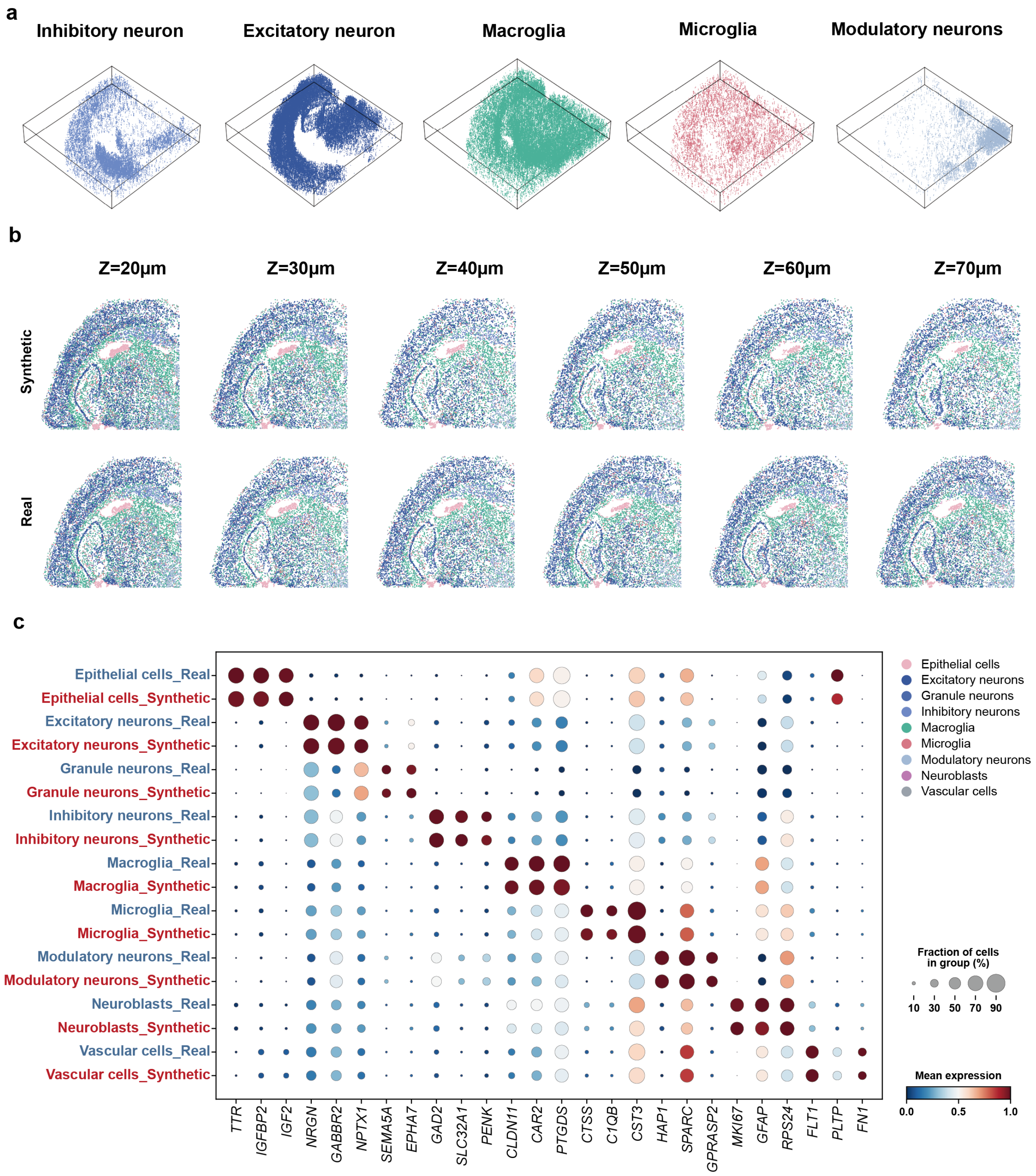


**Fig. S2.** Detailed spatial and transcriptomic characteristics of the 3D mouse brain Deep-STARmap dataset. **a,** 3D spatial distribution of distinct cell types in the mouse brain STARmap dataset. **b,** Virtual tissue slices generated by DEEPSPATIAL (synthetic) versus ground-truth real 3D tissue slices. **c,** Dot plot displaying gene expression patterns across different cell types.

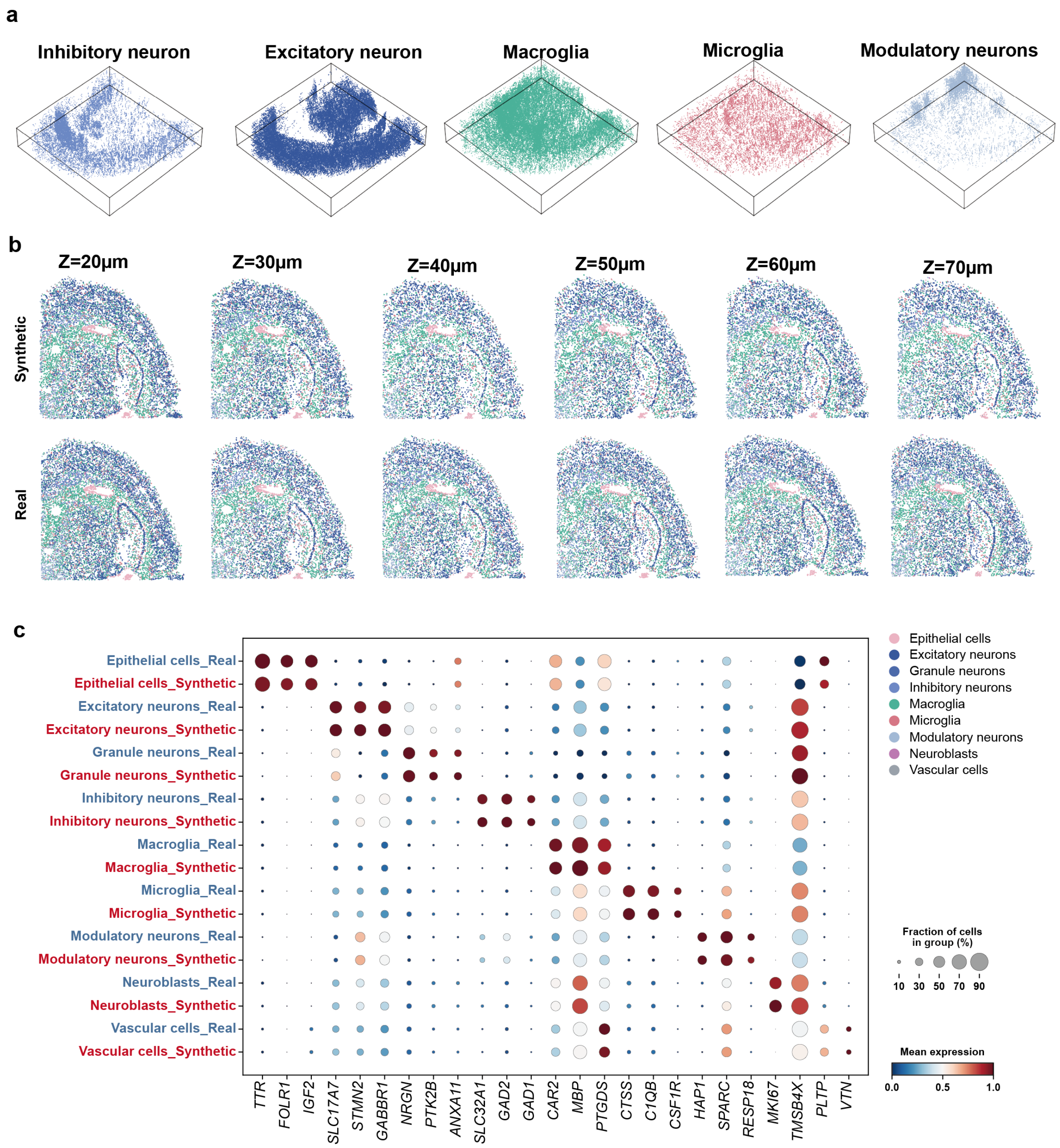


**Fig. S3.** Detailed spatial and transcriptomic characteristics of the 3D mouse brain Deep-RIBOmap dataset. **a,** 3D spatial distribution of distinct cell types in the mouse brain RIBOmap dataset. **b,** Virtual tissue slices generated by DEEPSPATIAL (synthetic) versus ground-truth real 3D tissue slices. **c,** Dot plot displaying gene expression patterns across different cell types.

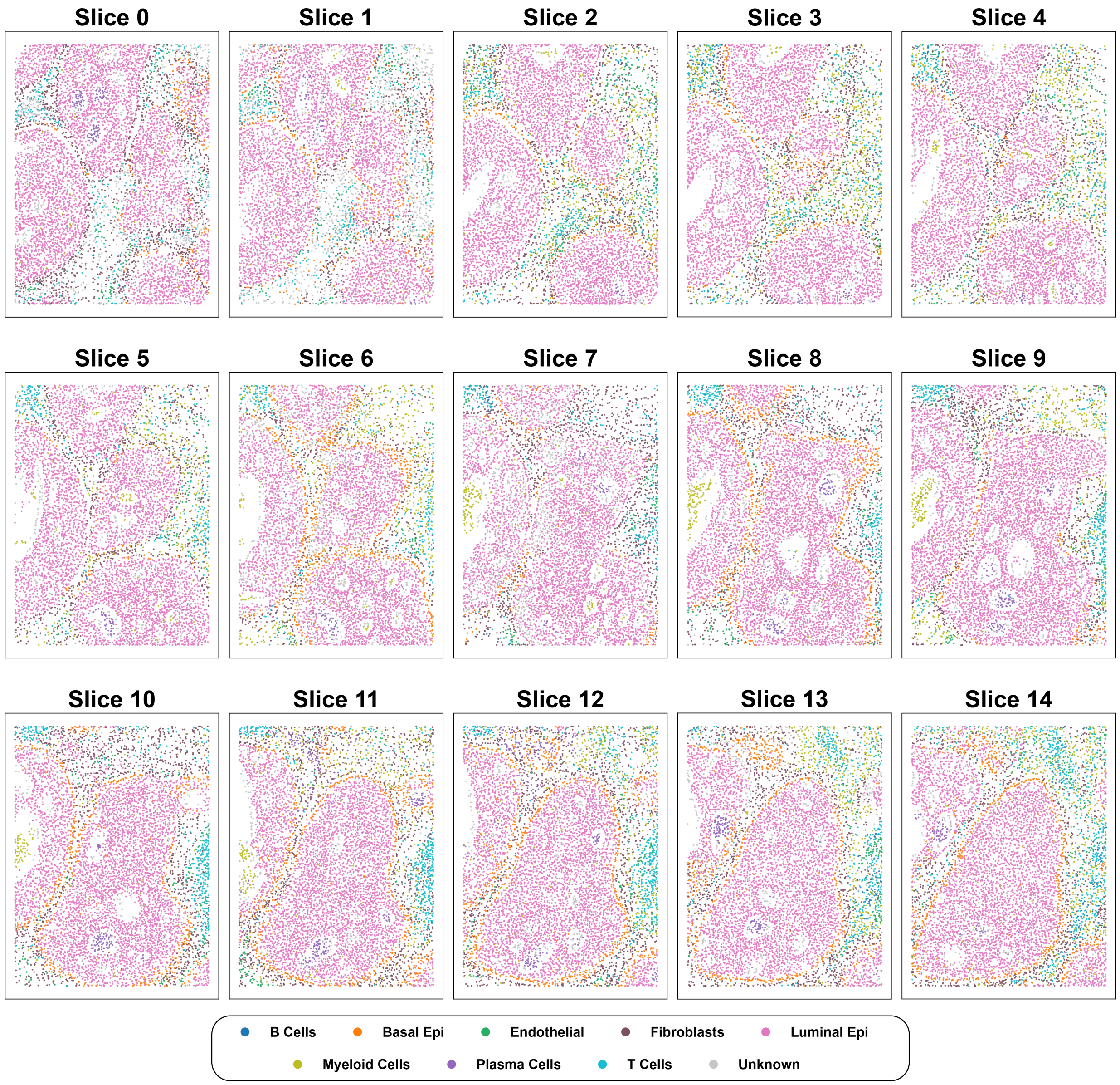


**Fig. S4.** Spatial distribution of cell populations across all experimental slices in the breast cancer (BRCA) IMC dataset.

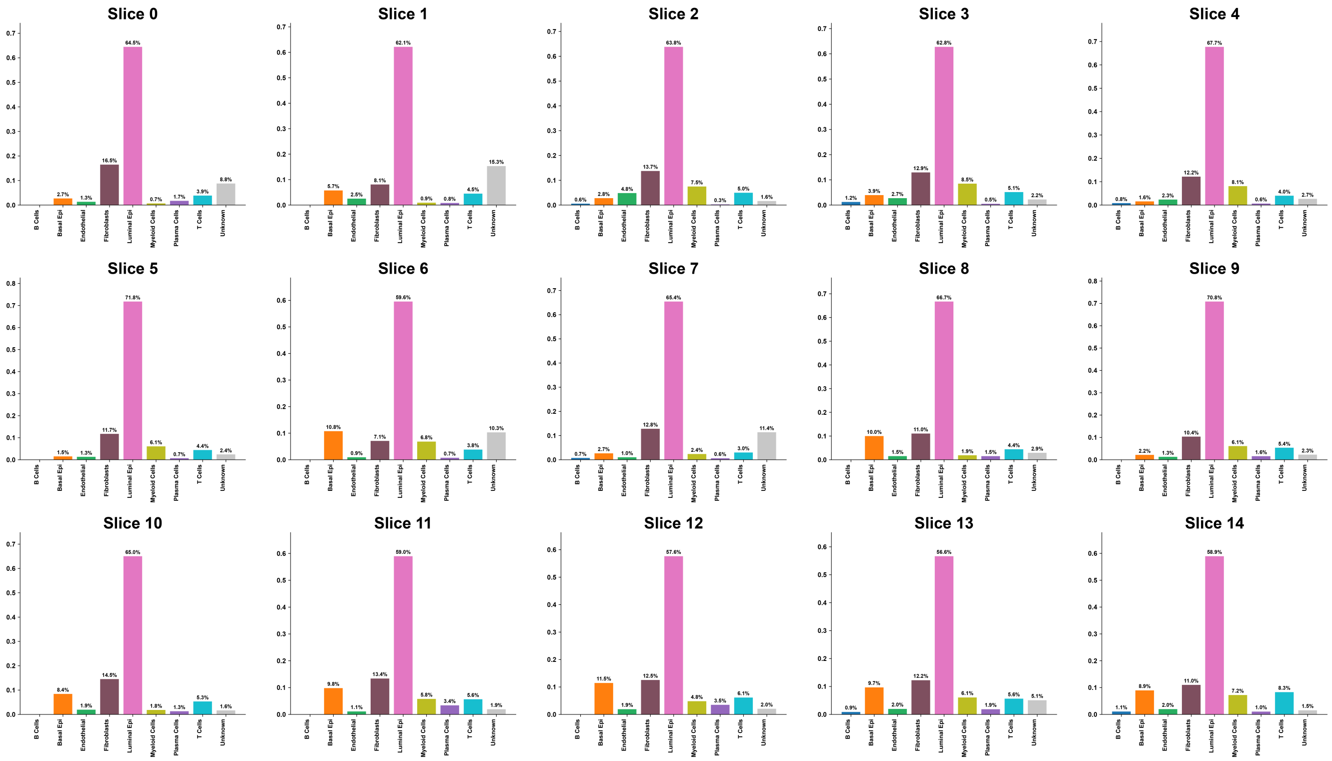


**Fig. S5.** Proportions of cell-type composition across all experimental slices in the BRCA IMC dataset.

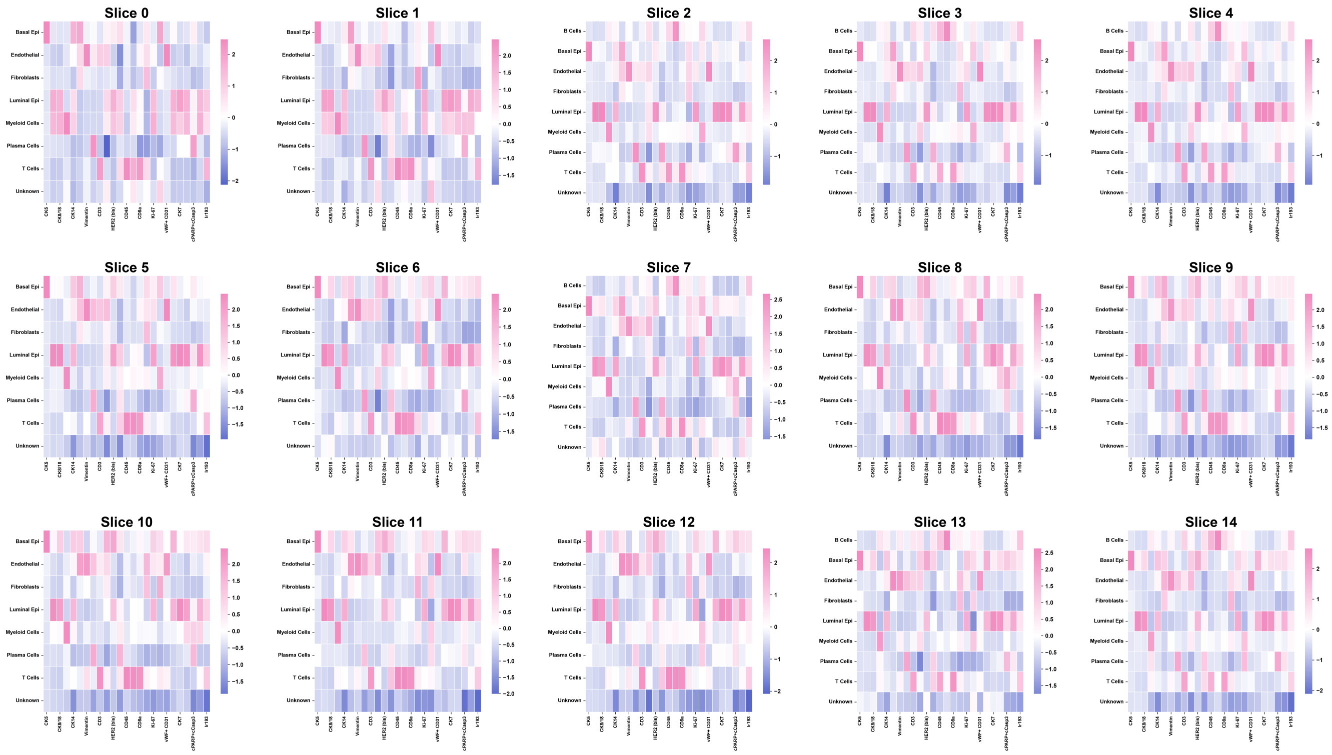


**Fig. S6.** Gene expression heatmaps across all experimental slices in the BRCA IMC dataset.

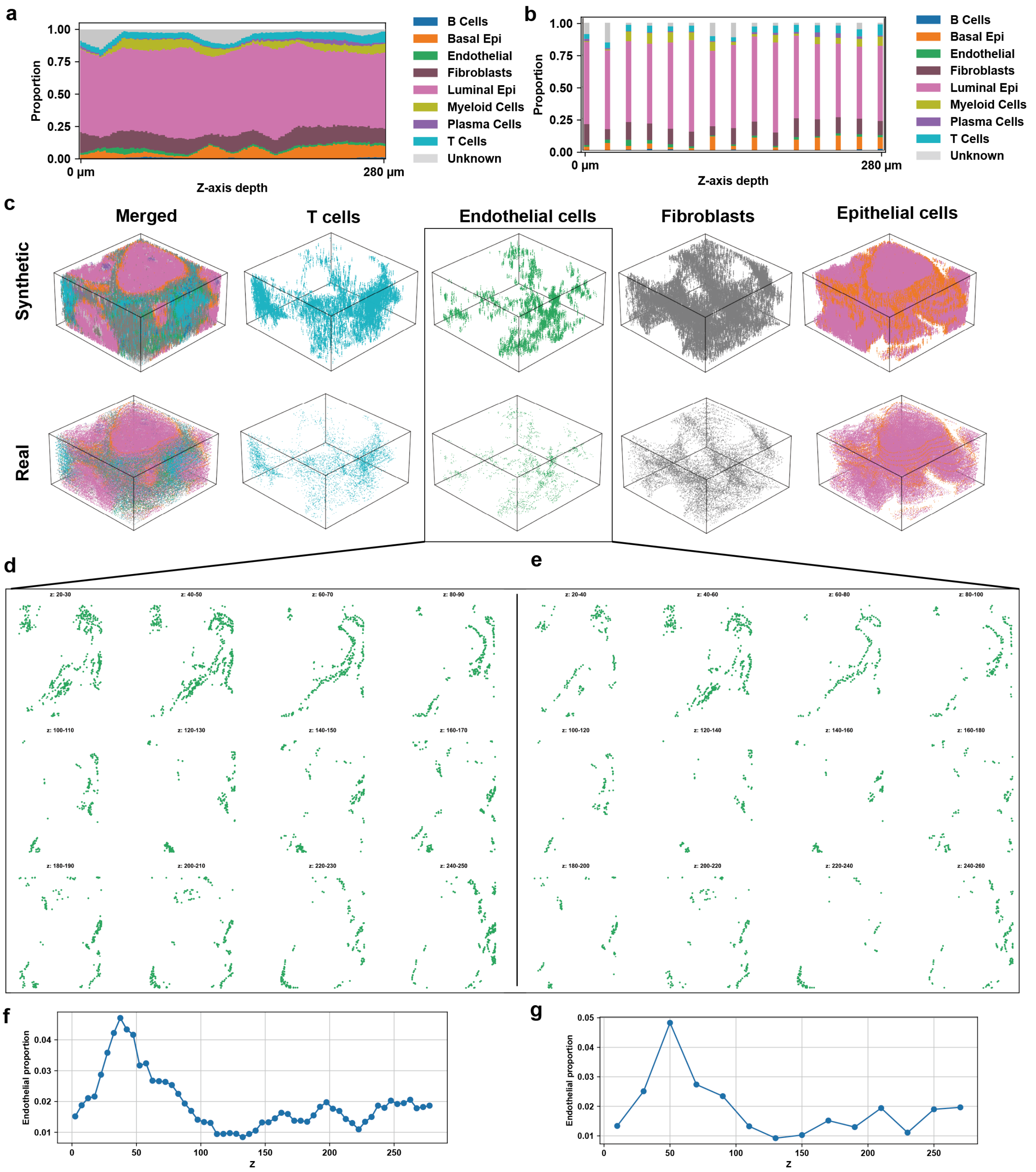


**Fig. S7.** Comprehensive comparison of vasculature-associated features between DEEPSPATIAL-reconstructed synthetic tissue and experimental BRCA IMC slices. **a,** Continuous cellular composition dynamics along the z-axis in DEEPSPATIAL-derived synthetic tissue. **b,** Discrete cellular composition observed in original experimental IMC slices. **c,** 3D spatial comparison of T cells, endothelial cells, fibroblasts and epithelial cells between synthetic and real tissue. **d,** Spatial distribution of endothelial cells in DEEPSPATIAL-generated virtual slices. **e,** Spatial distribution of endothelial cells in original experimental IMC slices. **f,** Z-axis variation in endothelial cell proportions in DEEPSPATIAL synthetic tissue. **g,** Z-axis variation in endothelial cell proportions in original experimental IMC slices.

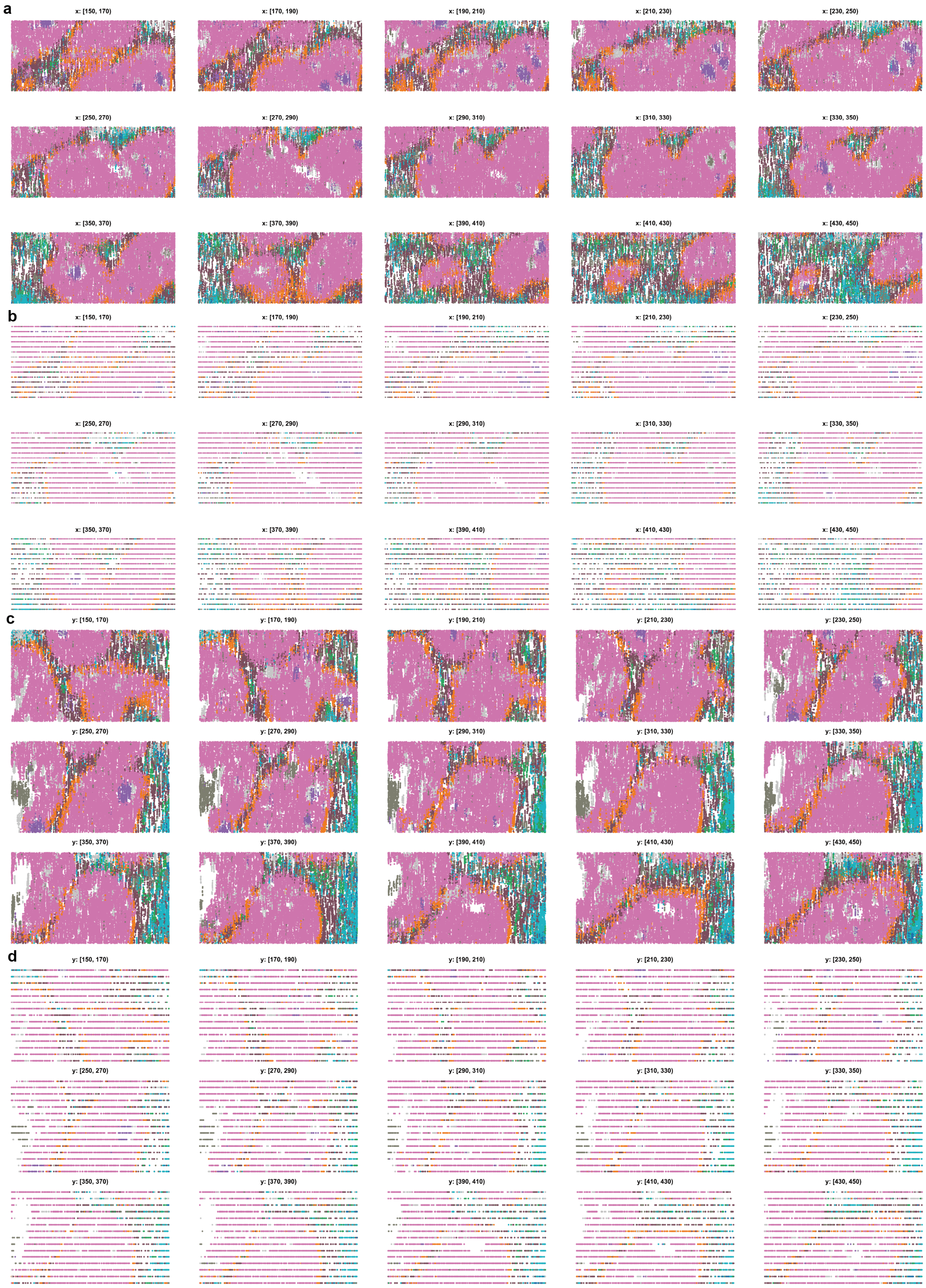


**Fig. S8.** Vertical virtual slice comparisons between DEEPSPATIAL-reconstructed synthetic tissue and experimental BRCA IMC data. **a,** YZ-plane virtual slices from DEEPSPATIAL synthetic tissue. **b,** YZ-plane slices from original experimental IMC data. **c,** XZ-plane virtual slices from DEEPSPATIAL synthetic tissue. **d,** XZ-plane slices from original experimental IMC data.

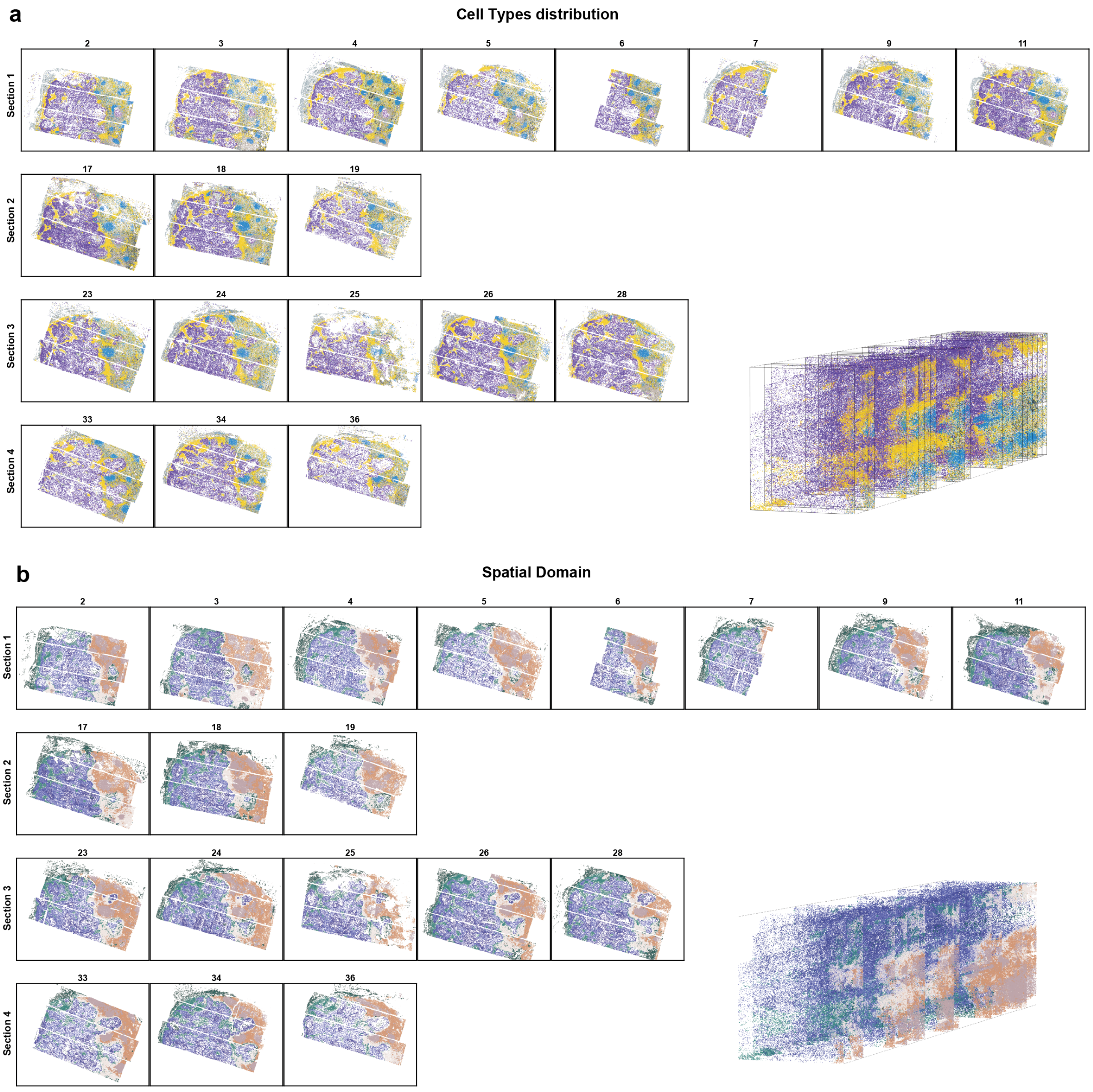


**Fig. S9.** Cellular spatial architecture of the openST HNSCC metastatic lymph node multi-slice dataset. **a,** Cell-type distribution across four representative experimental slices. **b,** Spatial domain distribution across four representative experimental slices.

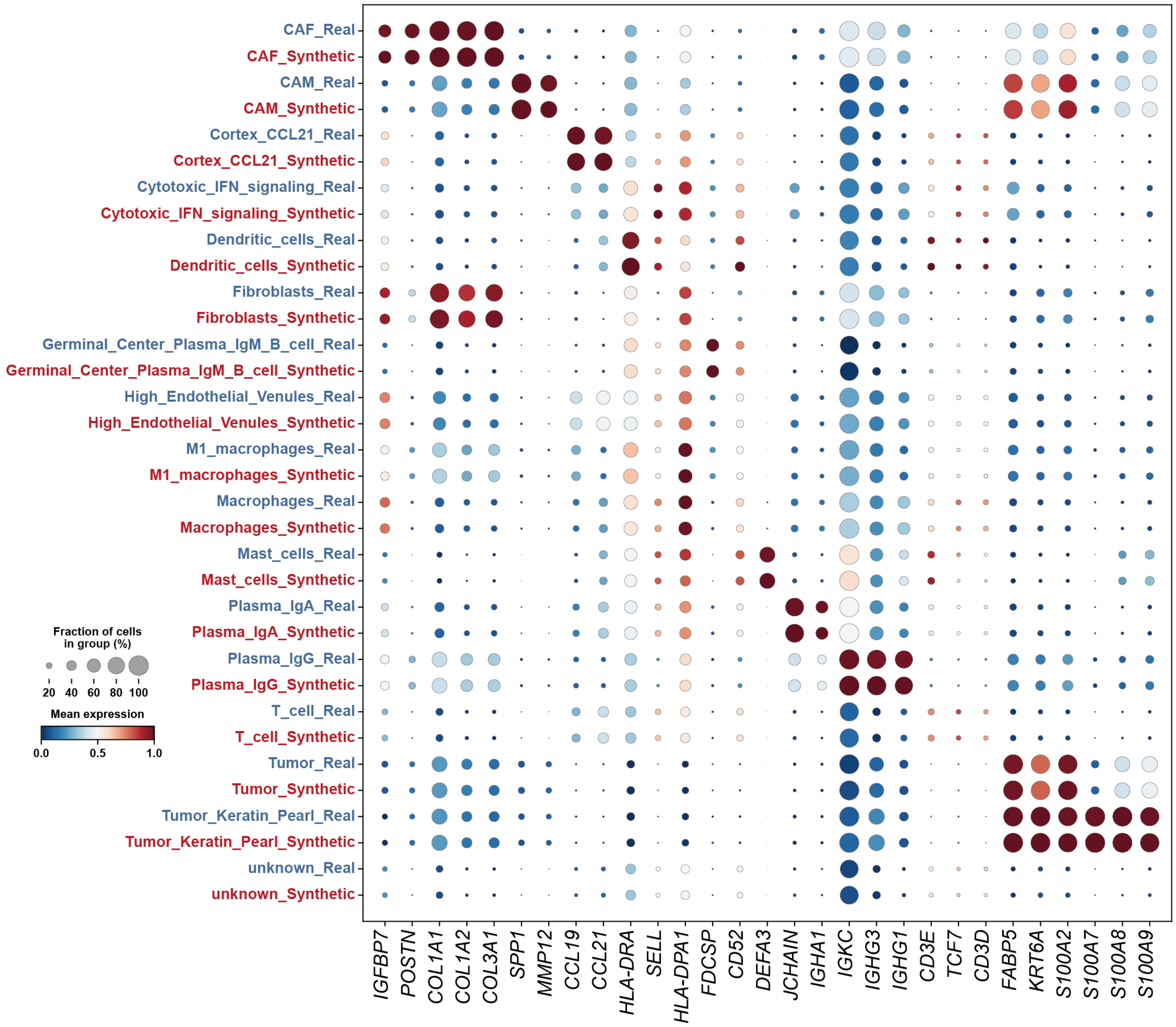


**Fig. S10.** Gene expression profiles of distinct cell types between DEEPSPATIAL-reconstructed synthetic data and original openST HNSCC dataset.

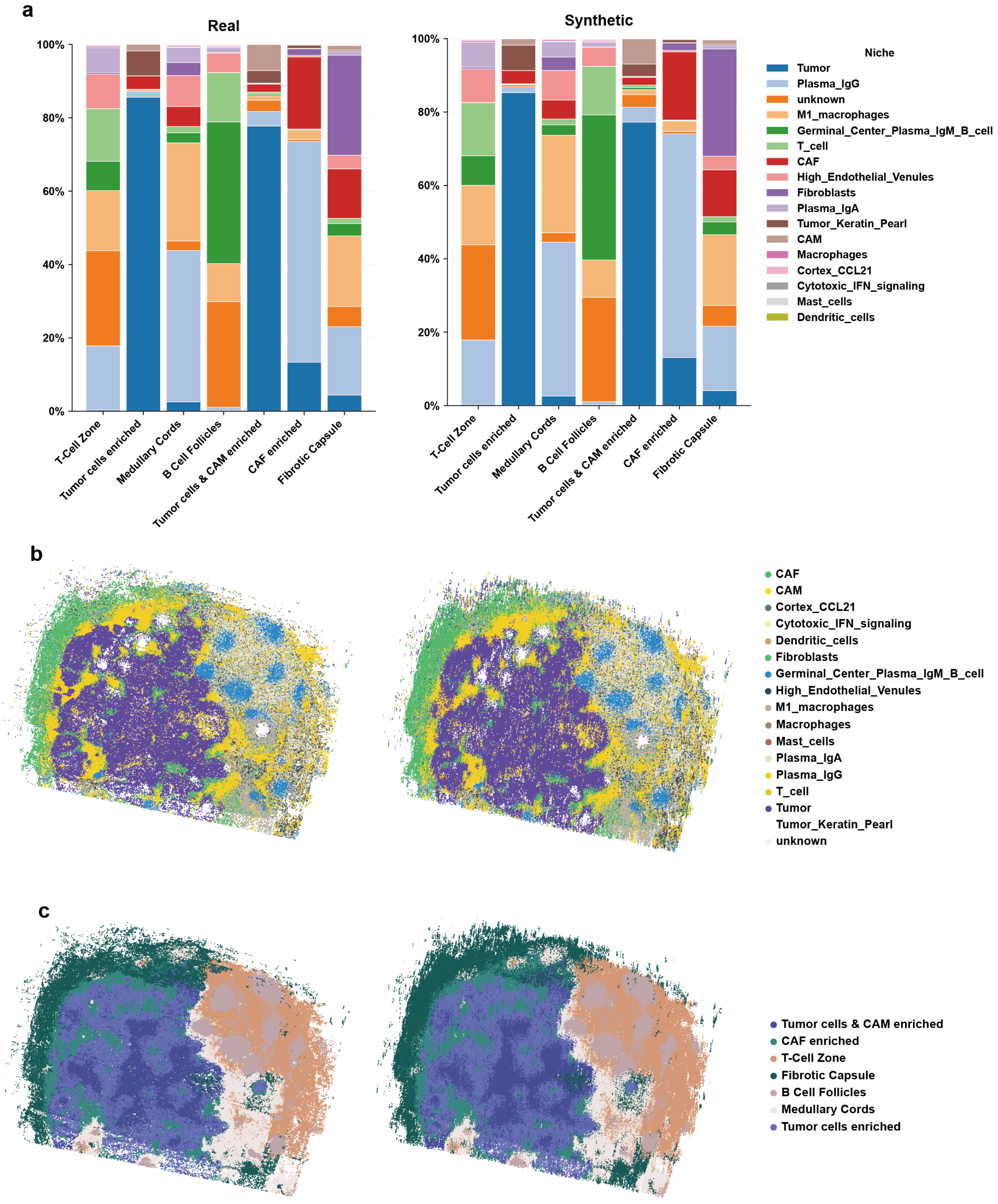


**Fig. S11.** Comparison of cell-type distribution within spatial domains between DEEPSPATIAL-reconstructed synthetic data and original openST HNSCC dataset.

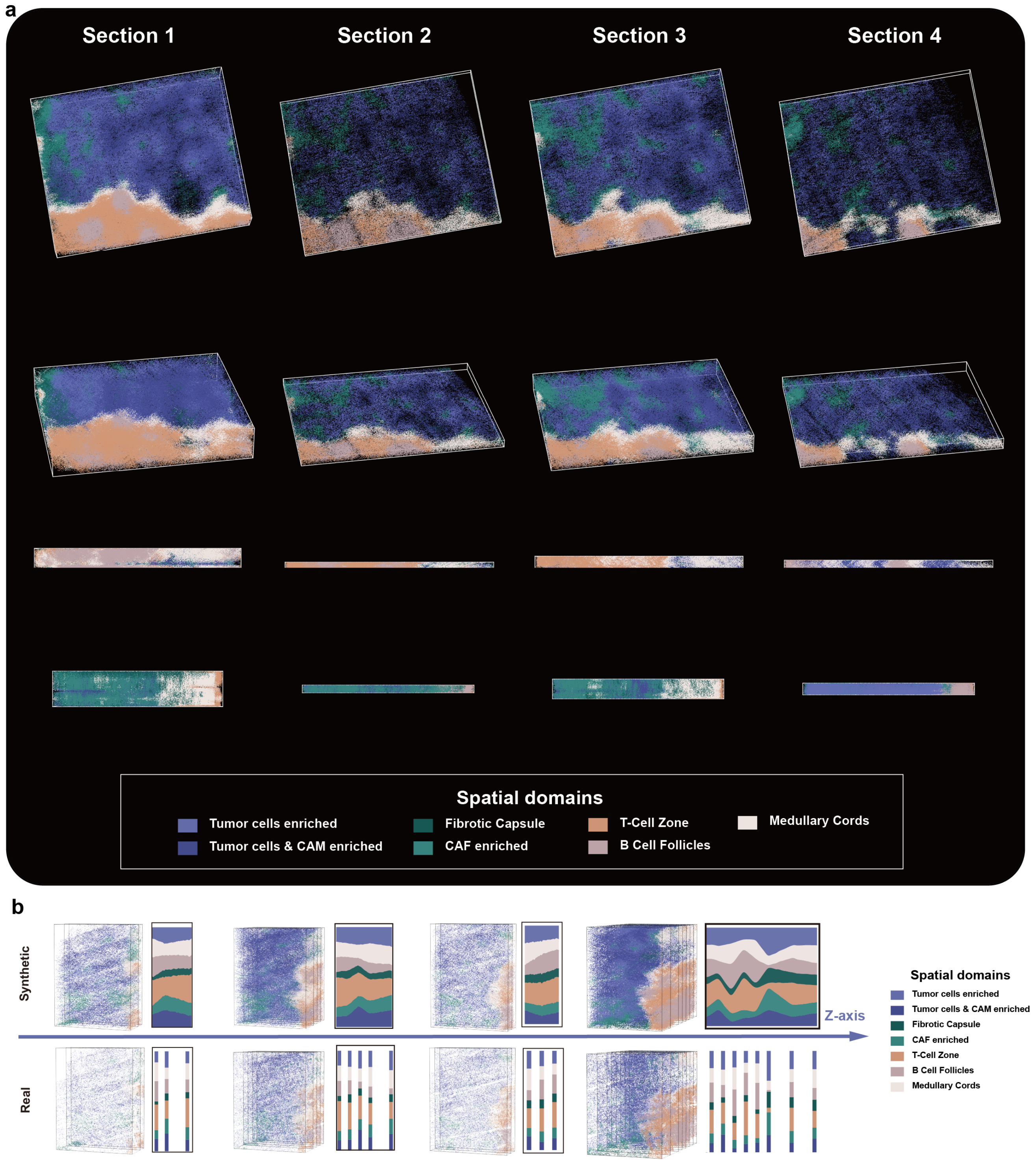


**Fig. S12.** 3D spatial domain architecture of the DEEPSPATIAL-reconstructed openST HNSCC dataset. **a,** 3D overview of spatial domain distribution. **b,** The spatial domain distribution through Z-axis.

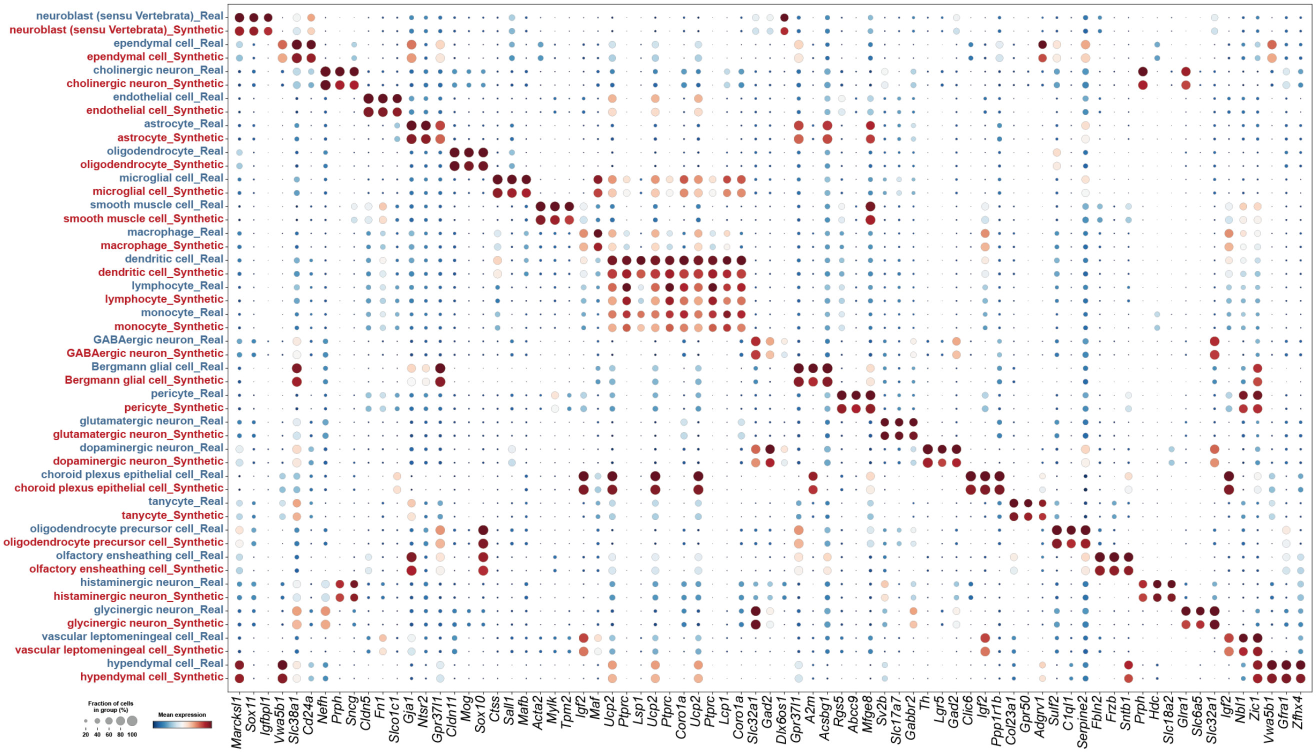

**Fig. S13.** Gene expression profiles of distinct cell types between DEEPSPATIAL-reconstructed synthetic data and the BRAIN Initiative Cell Census Network mouse brain dataset.

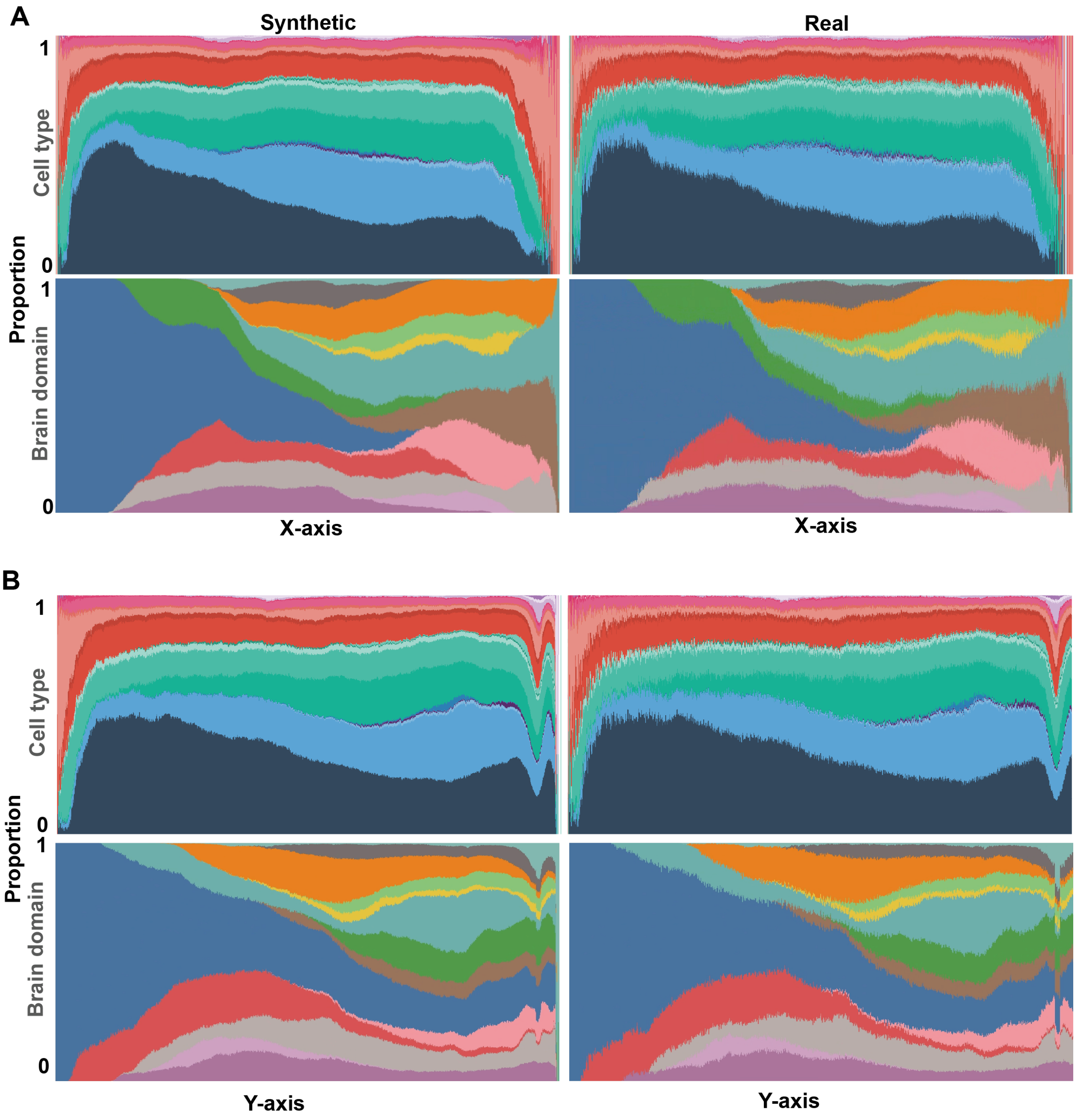


**Fig. S14.** Cell-type and brain region distribution along the X-axis (a) and Y-axis (b) between DEEPSPATIAL-reconstructed synthetic data and the BRAIN Initiative Cell Census Network mouse brain dataset.

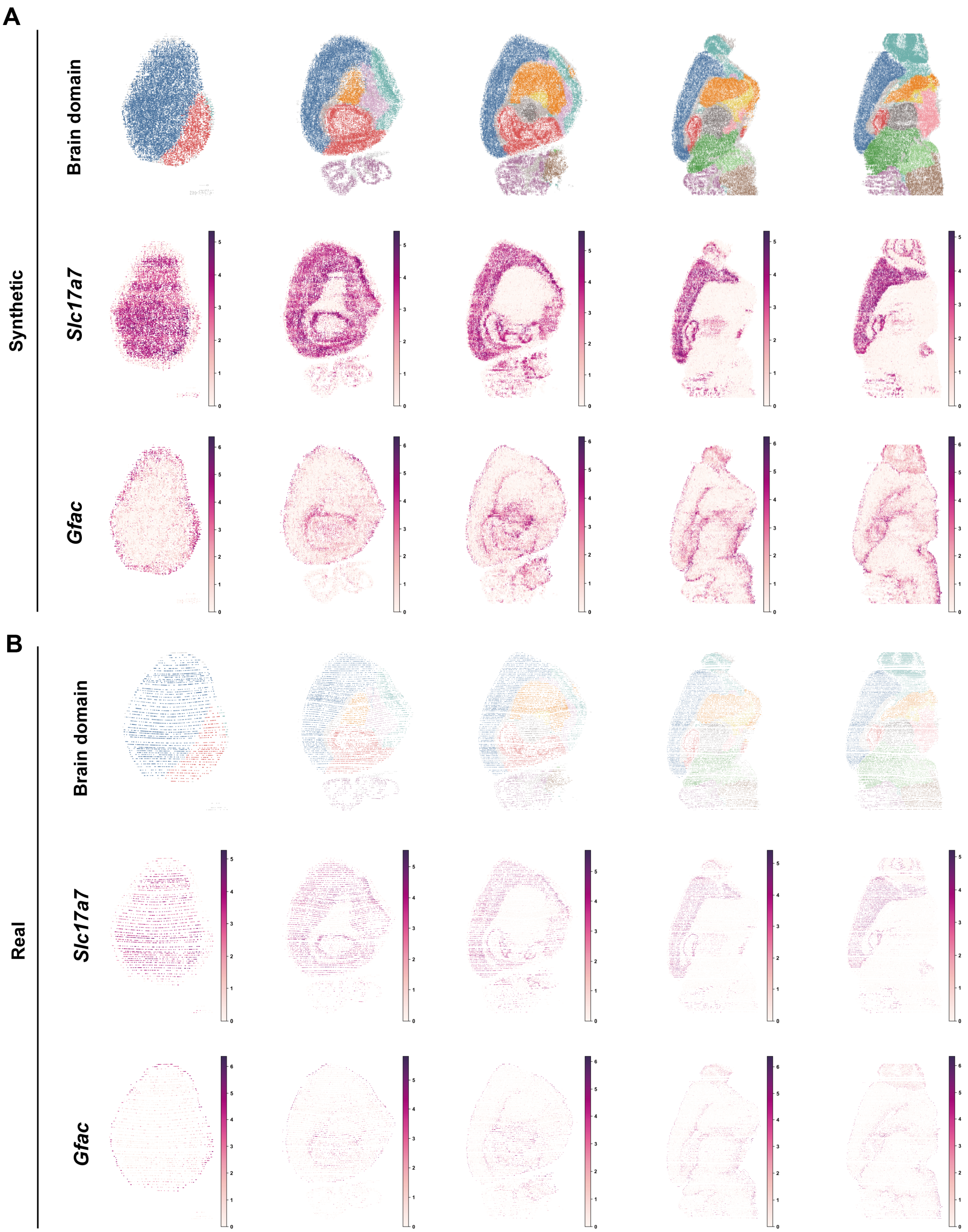


**Fig. S15.** Horizontal virtual slice comparison of coronal planes between DEEPSPATIAL-reconstructed synthetic data (a) and the BRAIN Initiative Cell Census Network real mouse brain data (b).

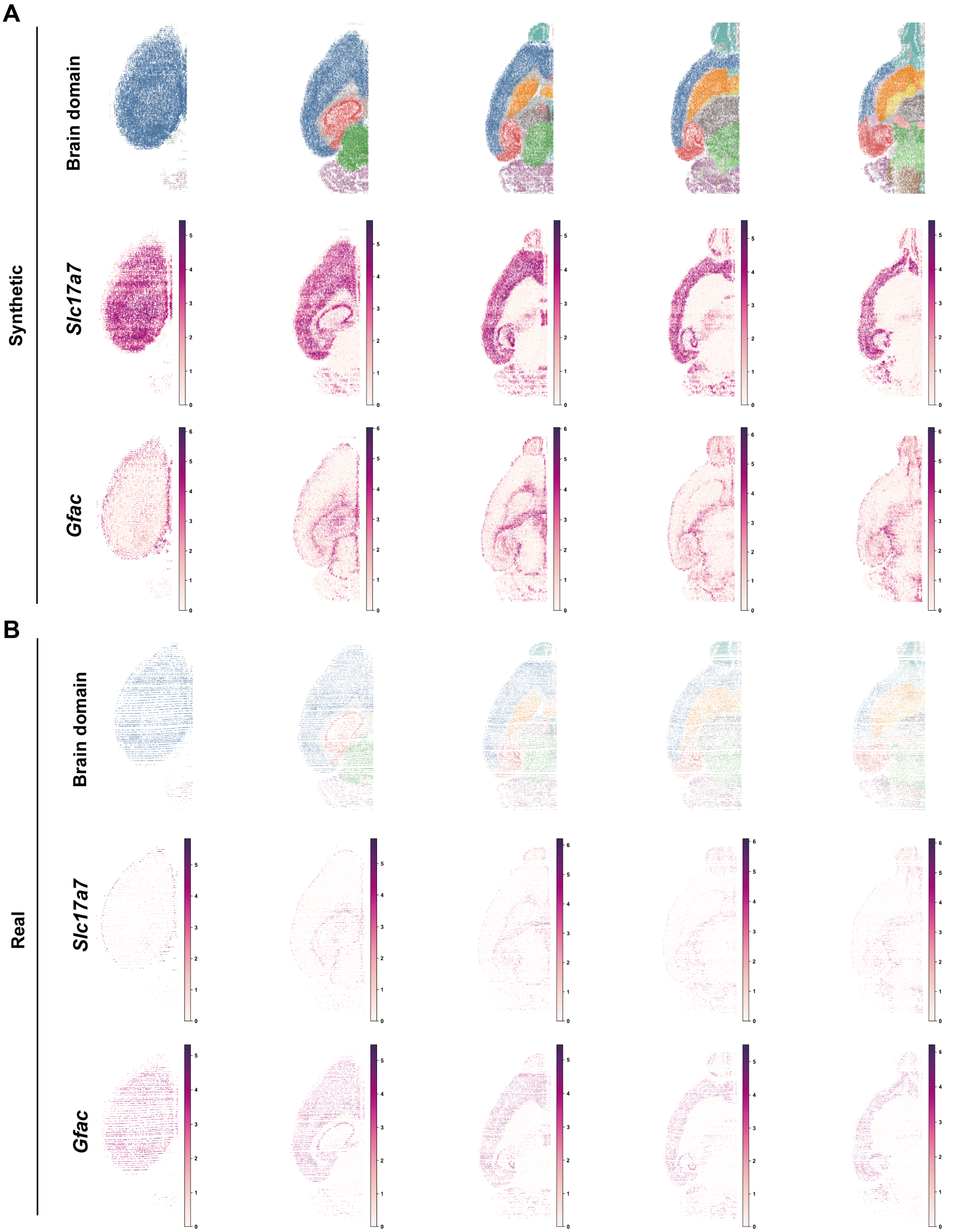


**Fig. S16.** Sagittal virtual slice comparison of coronal planes between DEEPSPATIAL-reconstructed synthetic data (a) and the BRAIN Initiative Cell Census Network real mouse brain data (b).
